# Particle algorithms for animal movement modelling in autonomous receiver networks

**DOI:** 10.1101/2024.09.16.613223

**Authors:** Edward Lavender, Andreas Scheidegger, Carlo Albert, Stanisław W. Biber, Janine Illian, James Thorburn, Sophie Smout, Helen Moor

**Affiliations:** Department of Systems Analysis, Integrated Assessment and Modelling, Eawag Swiss Federal Institute of Aquatic Science and Technology; School of Engineering Mathematics and Technology, University of Bristol; School of Mathematics and Statistics, University of Glasgow; School of Applied Sciences, Edinburgh Napier University; Centre for Conservation and Restoration Science, Edinburgh Napier University; Scottish Oceans Institute, University of St Andrews

**Author notes:** **Author contributions** Edward Lavender conceived the study and developed the methodology with Andreas Scheidegger, Carlo Albert and Helen Moor (Principal Investigator). This was motivated by work on the Movement Ecology of Flapper Skate project established by James Thorburn and an earlier modelling study led principally by Edward Lavender, Stanisław Biber, Janine Illian and Sophie Smout (Principal Investigator). In the current study, Edward Lavender led the analysis and writing of the manuscript, with substantive inputs from all authors. All authors contributed critically to the drafts and gave final approval for publication. **Data availability statement** Manuscript code is archived on Zenodo (Lavender et al., 2024a).

**Keywords:** animal tracking, flapper algorithm, movement ecology, passive acoustic telemetry, `patter`, utilisation distribution

## Abstract

1. Particle filters and smoothers are powerful sequential Monte Carlo algorithms used to fit non-linear, non-Gaussian state-space models. These algorithms are well placed to fit process-orientated models to animal-tracking data, especially in autonomous receiver networks, but to date they have received limited attention in the ecological literature.
2. Here, we introduce a Bayesian filtering–smoothing algorithm that reconstructs individual movements and patterns of space use from animal-tracking data, with a focus on passive acoustic telemetry systems. Within a sound probabilistic framework, the methodology uniquely integrates the movement process and the observation processes of disparate datasets, while correctly representing uncertainty. In a comprehensive simulation-based analysis, we compare the performance of our algorithm to the prevailing, heuristic methods used in passive acoustic telemetry systems and analyse algorithm sensitivity.
3. We find the particle smoothing methodology outperforms heuristic methods across the board. Particle-based maps consistently represent simulated movements more accurately, even in dense receiver networks, and are better suited to analyses of home ranges, residency and habitat preferences.
4. This study sets a new state-of-the-art for movement modelling in autonomous receiver networks. Particle algorithms provide a flexible and intuitive modelling framework with potential applications in many ecological settings.

## 1. Introduction

Animal movement shapes ecological processes across biological scales (Nathan et al., 2008, 2022). Individual movements reflect behaviour, such as foraging (Shaw, 2020), social interactions (Jacoby & Freeman, 2016) and habitat preferences (Abrahms et al., 2021). Individual movements also underlie emergent patterns of space use that shape population dynamics (Morales et al., 2010), ecosystem functioning (Riotte-Lambert & Matthiopoulos, 2020) and interactions with humans (Rutz et al., 2020).

Technologies for animal tracking have advanced dramatically in recent decades (Hussey et al., 2015; Kays et al., 2015; Nathan et al., 2022). Some technologies, such as satellite transmitters and global location sensors, record individual location (or proxies thereof) through time. Others, such as camera traps and acoustic telemetry, depend on autonomous receivers (often organised in static arrays) that record detections when animals move within range (Matley et al., 2022; Steenweg et al., 2017). These are cost-effective, long-term solutions for animal tracking, especially in forested ecosystems and underwater, where satellite tracking is limited (Hussey et al., 2015).

In aquatic ecosystems, passive acoustic telemetry is widely used to study individual movement across scales ranging from <10 km to >1000 km (Matley et al., 2022). This technology combines arrays of regularly or irregularly arranged acoustic receivers that listen continuously for individual-specific ‘pings’ from acoustic transmitters attached to animals. When animals move within a receiver’s detection range, detection(s) may be recorded (depending upon the distance between the receiver and the transmitter and other variables) (Kessel et al., 2014). Unfortunately, receiver detection ranges are typically non-overlapping, resulting in detection gaps when animal locations are less certain. However, additional devices (such as archival tags) may continue to collect data (such as depths) during this time (Matley et al., 2023).

Heuristic modelling approaches for individual movements and patterns of space use dominate in the passive acoustic telemetry community (Kraft et al., 2023). The predominant method (the ‘COA algorithm’) uses ‘centres of activity’ (COAs)—which are typically defined as weighted averages of the locations of receivers that recorded detections in successive time intervals—as ‘relocations’ for utilisation distribution (UD) estimation (Simpfendorfer et al., 2002; Udyawer et al., 2018). Other approaches, including network analysis (Lédée et al., 2015), refined shortest paths (RSP) (Niella et al., 2020) and the original ‘synthetic path’ methodology (Aspillaga et al., 2019), treat the receiver array as a network, with nodes defined by receivers and edges defined by sequential detections. For example, the RSP methodology uses interpolated ‘pseudo-relocations’ along the shortest paths between receivers (and distance-dependent errors) to fit a dynamic Brownian bridge movement model that represents the assumed pattern of space use. However, there remains little research on how well heuristic methods recover the underlying patterns of space use (i.e., method ‘performance’) and the circumstances in which they can be usefully applied (Lavender et al., 2023).

Recent studies have encouraged a shift from a heuristic to a process-orientated perspective that encapsulates the movement and measurement processes that generate observations (Hostetter & Royle, 2020; Lavender et al., 2023). State-space models formalise this perspective by coupling a process model of the unobserved movement process to an observation model of the measurement process that connects individual movements to observations (Patterson et al., 2008). Yet fitting state-space models is challenging. In the case of satellite telemetry, the Kalman filter is often used within a (linear) state-space modelling framework to refine location estimates and reconstruct trajectories, under the assumption that process and measurement errors are Gaussian (Jonsen et al., 2020). Particle filters generalise this methodology, providing a mathematically rigorous, flexible and intuitive modelling framework that is well-suited to movement modelling in autonomous receiver arrays.

The particle filter approximates the distribution of interest with a set of discrete samples termed ‘particles’ (Doucet & Johansen, 2009). In an animal-tracking context, the filter approximates the (partial) marginal distribution of the individual’s location (***s***) at time *t*, given all data (***y***) up to that time (that is, *f*(***s***_*t*_|***y***_1:*t*_)). The process model simulates particle movement over the landscape as a discrete-time Markovian walk (that is, ***s***_*t*_∼*f*(***s***_*t*_|***s***_*t*–1_)) and the observation model, together with a resampling step, eliminates or duplicates particles in line with the likelihood (that is, 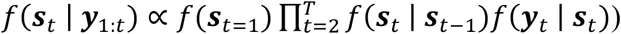. By including multiple observation models, it is straightforward to integrate disparate datasets that vary in quality and resolution. Subsequent smoothing algorithms approximate the full marginal distribution of the individual’s location at time *t*, given the data from the start (*t*=1) to the end (*t*=T) of the time series (that is, *f*(***s***_*t*_|***y***_1:*T*_)). Approximating the joint distribution (that is, sampling individual trajectories from *f*(***s***_1:*T*_|***y***_1:*T*_)) is also possible, but computationally much more expensive.

Particle filters have been widely used in engineering and physics (Doucet & Johansen, 2009) but only infrequently exploited in ecology (Andersen et al., 2007; Liu et al., 2019). In the field of acoustic telemetry, only one study that was principally framed in the context of existing ‘synthetic path’ methods has exploited the approach (Lavender et al., 2023). That study proposed a two-branch framework for reconstructing movements, comprising an acoustic-container (AC)-branch algorithm that resolved the set of possible locations for an individual, given the data, and a particle filter (PF)-branch algorithm that incorporated movement. Different combinations of AC-branch and PF-branch algorithms were collectively termed the ‘flapper algorithms’ and include the ACPF algorithm (which incorporates acoustic data) and the acoustic-container depth-contour (ACDC) PF algorithm (which incorporates acoustic and archival data). Here, we use the abbreviations ACPF and ACDCPF as generic labels for particle algorithms that incorporate acoustic or both acoustic and archival data (with or without smoothing).

In this study, we develop the use of particle algorithms for animal movement modelling in passive acoustic telemetry systems. Our methodology formalises and extends the ‘flapper algorithms’ within a powerful particle filtering—smoothing framework that (i) uniquely accounts for movement within and between periods of detection, (ii) realistically models the detection process and (iii) facilitates the integration of disparate datasets (such as acoustic and archival data). In a substantial simulation analysis, we validate the approach, analyse method performance and explore algorithm behaviour in different settings. The results robustly confirm our methods consistently outperform alternatives and represent a new state-of-the-art for movement modelling in passive acoustic telemetry systems.

## 2. Materials and methods

### 2.1. Overview

We formulate a Bayesian state-space model for the locations of a tagged animal in an acoustic telemetry system. Individual locations are denoted by ***s***. We consider the two-dimensional vector ***s*** = (*s*_*x*_, *s*_*y*_), where *s*_*x*_ and *s*_*y*_ are continuous. The movement process is represented as a regular series of steps from time *t*=1 to *t*=T, each of duration Δ*t*, by *f*(***s***_1:*T*_), where *f* is a probability density function. Contingent upon the individual’s location, observations (***y***), such as detections, are recorded at regular or irregular intervals. The objective is to infer the latent locations of the animal, using our knowledge of the movement process and the observations. Here, we formalise a model for this objective and a sampling algorithm. In the Supporting Information §1–5, we provide a complete algorithmic explanation for mathematical and non-mathematical readers. For a notational summary, see Table S1.

### 2.2. Model

#### 2.2.1. Objective

Formally, our objective is to derive the joint probability distribution *f*(***s***_1:*T*_|***y***_1:*T*_) of a tagged individual’s possible trajectories from the start to the end of the time series, accounting for the movement process and the observations. Using Bayes’ theorem, the joint distribution can be expressed in terms of a prior (the movement process) and a likelihood (the observation process), i.e.,

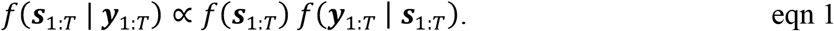

#### 2.2.2. Movement process

The expression *f*(***s***_1:*T*_) represents movement. We model *f*(***s***_1:*T*_) as a discrete-time Markovian process

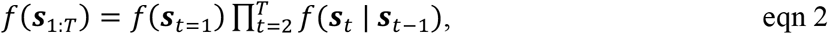

where *f*(***s***_*t*=1_) is a probability density function of the individual’s starting location and *f*(***s***_*t*_|***s***_*t*–1_) is the probability density of moving from ***s***_*t*–1_ → ***s***_*t*_, under the assumption that an individual’s location at each time step depends only on its previous location. A simple model for *f*(***s***_*t*_|***s***_*t*–1_) is a restricted two-dimensional random walk, in which the location ***s***_*t*_ is given by

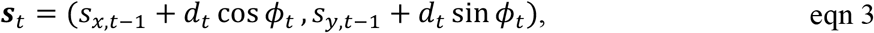

where *d* and *ϕ* are independently distributed random variables that represent step lengths and turning angles, subject to boundary conditions (i.e., land). In this paper, we use a Gamma step-length distribution based on pre-defined shape (*k*) and scale (*θ*) parameters and a truncation interval defined between zero and mobility:

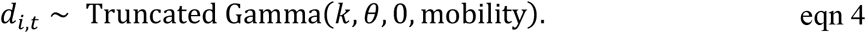

For turning angles, we use a uniform distribution:

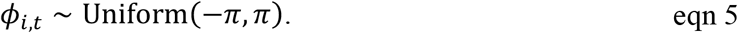

#### 2.2.3. Observation process

##### 2.2.3.1. Joint likelihood

The term *f*(***y***_1:*T*_|***s***_1:*T*_) denotes the joint likelihood. This measures the probability of the observations given the locations of the individual (***s***_1:*T*_). Under the assumption that the observations, conditional on ***s***_1:*T*_, are independent, we express *f*(***y***_1:*T*_|***s***_1:*T*_) as the product of the likelihood from each time step:

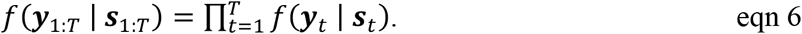

Observations may include acoustic records and ancillary data, such as depth measurements. Acoustic observations comprise detections, which are explicitly recorded, and non-detections, which are implicitly known for each *t* (assuming Δ*t* exceeds the transmission interval). We use ***y***^(*A*)^ to denote an *M* × *T* matrix of acoustic observations, where *M* is the number of receivers, and ***y***^(*D*)^ to denote a row vector of depth observations (from 1, . ., *T*). Both sets of observations form the dataset ***y*** = {***y***^(*A*)^, ***y***^(*D*)^}. The combined likelihood at time *t* is given by

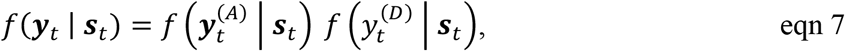

where 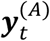 is a column vector of acoustic observations at time *t* and 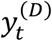 is the depth observation.

##### 2.2.3. Acoustic observations

The acoustic likelihood expresses the correspondence between acoustic observations and latent locations. We denote a detection at receiver *k* as 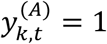 and non-detection as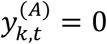. We assume detections are the outcomes of a Bernoulli process and express the probability of the observation 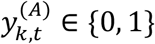 at receiver *k* as:

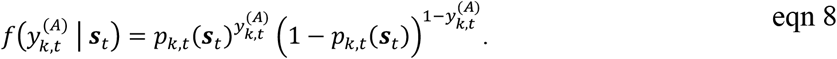

Detection probability (*p*) is modelled as a function of covariates. A simple model for *p*, of the kind commonly estimated from range-testing data, is a logistic function of the distance between the receiver’s location (***r***_1_ = (*s*_*x*_, *s*_*y*_)) and transmitter (***s***), that is

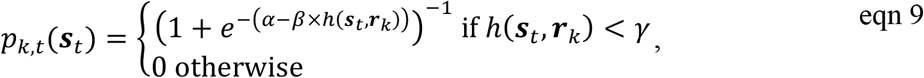

where *h*(⋅,⋅) is a distance function; *α* and *β* are parameters; and *γ* is the detection range (Lavender et al., 2023).

The combined probability of all acoustic observations is the product of independent probabilities from each receiver:

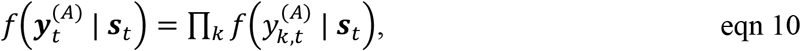

where *k* indexes over operational receivers at time *t*.

##### 2.2.3.3. Depth observations

The depth likelihood expresses the correspondence between the depth observation 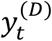 at time *t* and the latent location. An example model for the probability density of 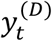 is

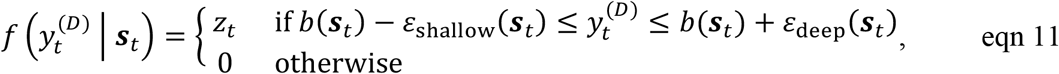

where *z*_*t*_ = ((*ε*_deep_ (***s***_*t*_) + *ε*_*s*hallow_ (***s***_*t*_))^−1^. This model requires 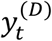 to be within an envelope around the bathymetric depth *b*(***s***_*t*_) defined by the shallow- and deep depth-adjustment functions, *ε*_shallow_ (***s***_*t*_), *ε*_deep_ (***s***_*t*_) ≤ *b*(***s***_*t*_). These functions capture observational uncertainty and spatially explicit bathymetric uncertainty and can be tailored to species with different lifestyles: for benthic species, observations must be close to the seabed, which can be enforced by small errors (*ε*_shallow_ (***s***_*t*_), *ε*_deep_ (***s***_*t*_) ≪ *b*(***s***_*t*_)); for pelagic species, observations may occur in the water column, which is permitted by larger *ε*_shallow_ (***s***_*t*_) values.

### 2.3. Sampling algorithm

#### 2.3.1. Particle filter

We have formulated a model for the joint distribution of the individual’s locations, given all data (that is, *f*(***s***_1:*T*_|***y***_1:*T*_)). We now outline a sampling algorithm. The target of our inference is location; we assume static parameters are known. For inference, we begin with the simpler, partial marginal distribution for the possible locations for an individual at time *t*, given the data up to that time and the movement model (that is, *f*(***s***_*t*_|***y***_1:*t*_)). This is iteratively represented as

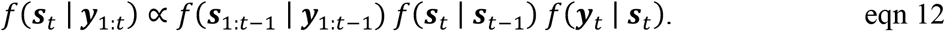

The particle filter approximates *f*(***s***_*t*_|***y***_1:*t*_) as a sum of *N* weighted samples, termed ‘particles’, that is,

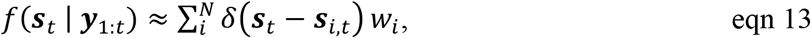

where *δ* is the Dirac delta function, *w* denotes normalised weights (∑_*i*_ *w*_*i*_=1) and *i* indexes particles, which represent possible locations for the individual. This is an iterative procedure (derived from eqn 12) that at each time step comprises three stages:

- **Simulation (movement)**. We simulate particles, following eqns 2–5. Initial particles are sampled from a probability distribution, such as a uniform distribution, via ***s***_*i,t*=1_ ∼*f*(***s***_*t*=1_). At subsequent time steps, we simulate particles from the movement process, that is ***s***_*i,t*_ ∼*f*(***s***_*i,t*_ | ***s***_*i,t*–1_), using a movement model, such as a random walk (eqn 3).
- **Weighting (observation)**. Particles are weighed by their compatibility with the observations, in line with the likelihood, that is *w*_*i,t*_ ∝ *w*_*i,t*–1_*f*(***y***_*t*_|***s***_*i,t*_), following eqns 6–11.
- **Resampling**. Periodically, particles are re-sampled in line with the weights. This procedure eliminates unlikely particles and duplicates likely ones.

The time complexity of the particle filter is 𝒪(*NT*) (Doucet & Johansen, 2009).

The end result is a set of particles (location samples) from *f*(***s***_*t*_|***y***_1:*t*_) that represent the possible locations of the individual at each time step, given all preceding and contemporary data. Theoretically, a small number of particles (*N* ≈ 1000) is sufficient to approximate a two-dimensional distribution, of the kind described here, but in practice larger numbers of particles may be required in the filter to ensure that sufficient particles remain ‘alive’ at each time step to approximate *f*(***s***_*t*_|***y***_1:*t*_); that is, to achieve convergence.

#### 2.3.2. Particle smoothing

Particle smoothing iteratively re-weights particles from the particle filter to approximate the full marginal, *f*(***s***_*t*_|***y***_1:*T*_) (Doucet & Johansen, 2009). The two-filter smoother uses *N* particles (***s***_*i,t*_) from a forward run of the particle filter (with weights *w*_*i,t*_) and *N* particles 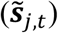 from a backward run (with weights 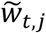) to obtain a set of smoothing weights 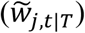:

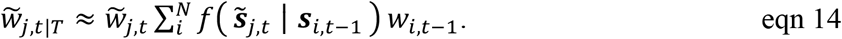

For each particle 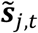, these weights effectively sum over all possible movements from the preceding particles on the forward filter. The distribution *f*(***s***_*t*_|***y***_1:*T*_) is approximated as a weighted sum of smoothed particles:

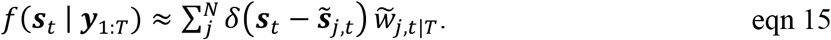

The time complexity of smoothing is 𝒪(*N*^2^*T*) (Doucet & Johansen, 2009). However, typically only a subset of the particles required for a successful run of the particle filter are required for an effective approximation of *f*(***s***_*t*_|***y***_1:*T*_).

### 2.4. Mapping

Particles can be used to reconstruct movements and map patterns of space use. For mapping, we suggest the ‘probability-of-use’ metric (*P*), which represents the probability that an individual is located in a given location at a randomly chosen time. This can be calculated across a grid as weighted average of the particles in each cell:

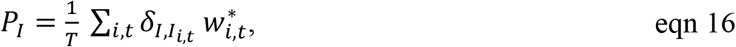

where *I* indexes grid cells, *I*_*i,t*_ is the grid cell of particle *i* at time *t*, 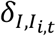 is the Kronecker delta and 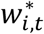 is the weight of that particle at time *t* (either *w*_*i,t*_, 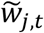 or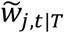). Particles may be derived from the filter (approximating *f*(***s***_*t*_|***y***_1:*t*_)) or the smoother (approximating *f*(***s***_*t*_|***y***_1:*T*_)), which is more expensive but produces refined maps of space use. In practice, *P*_*I*_ is sensitive to grid resolution and we suggest computing *P*_*I*_ across a fine grid followed by kernel smoothing (see Supporting Information §6).

### 2.5. Simulations

#### 2.5.1. Software

We developed the `patter R` package (Lavender, 2024a) and the `Patter.jl` backend (Lavender, 2024b) to implement the methodology (Lavender et al., 2024b). Here we illustrate and evaluate algorithm performance and sensitivity by simulation, using R, v.4.3.1 (R Core Team, 2023). Code is available online (Lavender et al., 2024a).

#### 2.5.2. Study systems

In all simulations, we considered a ∼100 km^2^ rectangular area and a 10 × 10 m bathymetric grid. Within this area, we considered two hypothetical ‘study systems’ defined by distinct movement and observational processes, for a hypothetical benthic animal and acoustic and archival data (Fig. S1, Table S2). (Two study systems were considered to validate the robustness of our simulations to the parameters characterising any one system.) Both movement processes were defined by a discrete-time, continuous-space Markovian random-walk model (eqns 2–5), with the individual taking a ‘step’ up to ‘mobility’ metres in length every 2 min, over a 2-day period (Table S2). The detection process was binomial, with detection probability declining logistically to zero by *γ* metres from receiver(s), following eqn 9 (Table S2). The archival observation process was uniform and constant across all simulations (Table S2). In each system, we simulated 20 acoustic arrays with 10, 20, …, 100 receivers, arranged randomly or regularly (Fig. S2, Table S3). In each system and for each array, we simulated 30 realisations of the movement process (30 movement paths) and 30 corresponding realisations of the observation processes (30 acoustic and archival datasets). We only considered the simulated paths and observations between the first and last detection and simulations which generated > 10 detections.

#### 2.5.3. Performance

We compared patterns of space use exhibited by simulated paths to those reconstructed from the simulated observations by the COA, RSP and particle algorithms (ACPF and ACDCPF). For simulated paths, ‘true’ patterns of space use were generated by fitting kernel UDs to path coordinates using cross validation. These UDs were compared against the UDs generated from each algorithm, visually and with standard error metrics, such as the mean error (the mean difference between a ‘true’ and reconstructed UD, denoted ME). For details, see Supporting Information §7–8 and Table S4.

#### 2.3.4. Sensitivity

For a subset of arrays, we analysed particle algorithm sensitivity. The purpose of this analysis was to investigate how patterns of space use change if the parameters of the process models (such as γ), which are assumed known, are overly restrictive or flexible. To investigate algorithm sensitivity, we compared UDs for simulated paths to those reconstructed by particle filtering–smoothing algorithm implementations with mis-specified parameters (for *k, θ*, mobility, *α, β* and *γ*), qualitatively and with ME (Fig. S3, Table S5). This analysis reveals the extent to which patterns of space use are sensitive to selected parameters and which parameters should therefore be prioritised in data-collection efforts (such as range tests).

## 3. Results

### 3.3. Performance

In the visual analysis of UDs, heuristic methods were consistently outperformed by particle algorithms (Figs 1 and S4). In the sparse arrays (with 10 receivers: Fig. 1A–L), COAs concentrated in specific areas and poorly represented underlying patterns, irrespective of receiver arrangement (e.g., Fig. 1A versus 1B). RSP maps also misplaced hotspots, concentrating them around receivers, but by smoothing the connections between receivers better represented those transitions and exhibited lower ME (e.g., Fig. 1C). The particle algorithms suggested more nuanced patterns (e.g., Fig. 1D–E). Maps based on the particles from the forward filter (approximating *f*(***s***_*t*_|***y***_1:*t*_)) were relatively diffuse, but more accurately placed hotspots away from receivers, resulting in an ME similar to RSPs (Fig. S4). Particles from the smoother (approximating *f*(***s***_*t*_|***y***_1:*T*_)) produced more refined maps, with lower ME than RSPs in both random and regular arrays (Fig. 1D–F and 1J–L). Compared to the ACPF algorithm, the ACDCPF algorithm suggested more restricted patterns of space use, in line with the contours of the bathymetric landscape that we simulated (Figs 1E, 1K, S2 and S4).

**Fig. 1.**
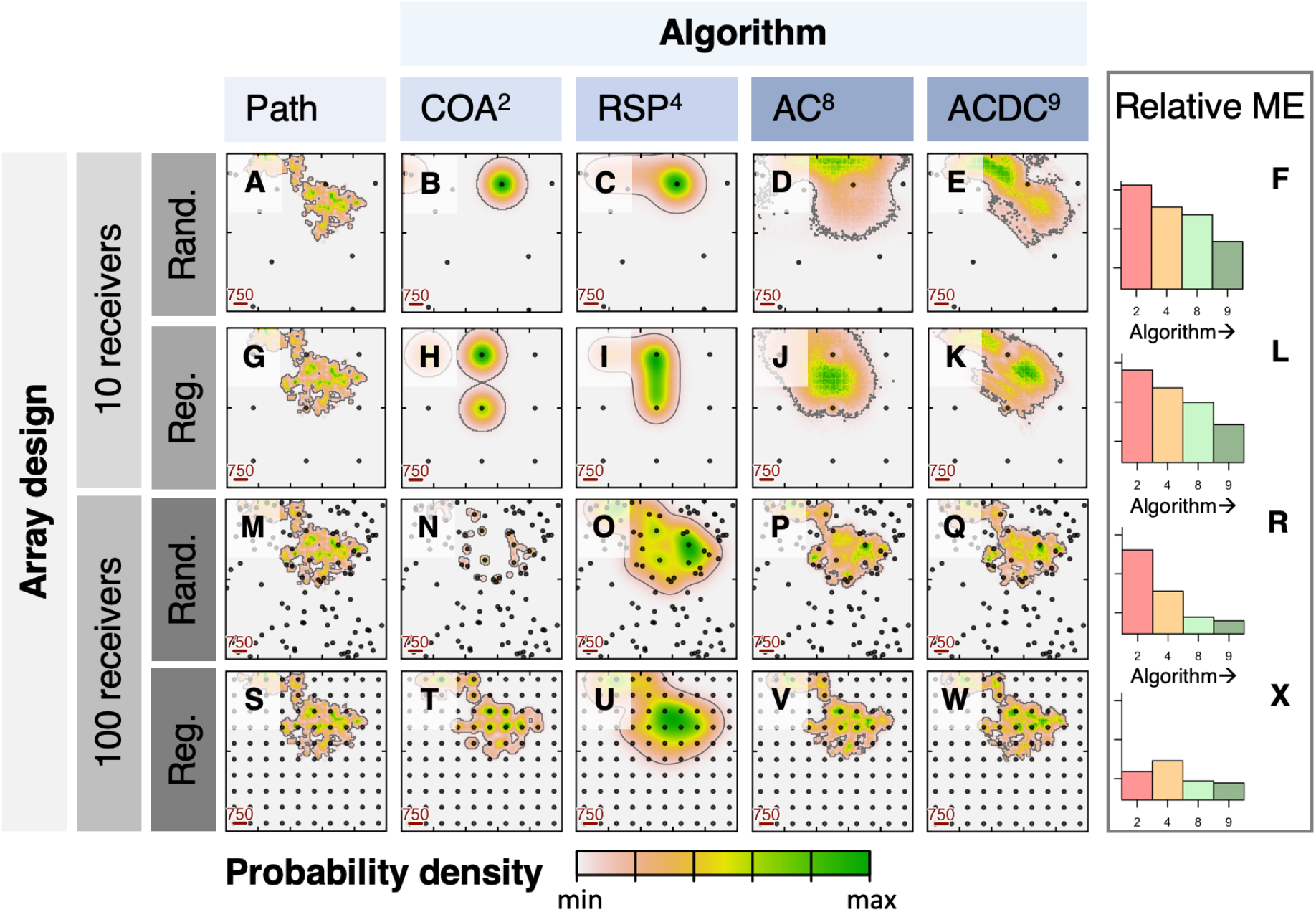
Utilisation distributions (UDs) reconstructed by different algorithms in selected array designs for a hypothetical study system. Four designs with 10 or 100 receivers in random or regular arrangements are shown. For each array (row), the estimated UD for the portion of the simulated path between the first and last detections is shown (hence slight differences between rows), alongside comparable UDs generated by the COA, RSP and particle filtering–smoothing algorithms (ACPF and ACDCPF). Lines mark home ranges (and contain 95 = of the probability density volume). Points mark receivers. The detection range was 750 m. Bar plots show relative mean error (ME). Algorithm superscripts identify selected algorithm implementations (of which there were nine in total), following Table S4. Panels correspond to the first simulated study system. For the full figure, see Fig. S4.

In the dense arrays (with 100 receivers), algorithms represented underlying patterns more effectively (Figs 1M–X and S4). For the COA algorithm, there was a clear influence of receiver arrangement, with heavy fragmentation (and high ME) in the random array (Fig. 1N) and smoother patterns (and lower ME) in the regular array (Fig. 1T). RSPs behaved similarly in the two array designs, broadly capturing but over-smoothing simulated movements (Fig. 1O and 1U). Accordingly, the COA maps were worse than RSP maps in the random array (Fig. 1R) but marginally better in the regular array according to ME (Fig. 1X). The particle algorithms performed more effectively in both array designs, producing accurate maps that closely correspond to those for the simulated path (Fig. 1P–R and 1V–X). Differences between maps from the filter and the smoother were scarcely apparent (Fig. S4).

In the illustrated arrays (with 10 or 100 receivers), a consistent ranking of algorithms emerged from repeated simulations of the same data-generating processes (Figs 2 and S5). In the sparse arrays (with 10 receivers), the COA algorithm generally produced the highest ME (occasionally even exceeding a null model) (Fig. 2A–B). RSPs and the particle algorithms produced overlapping, but consistently lower, MEs (Fig. 2A–B). In general, the forward particle filter produced marginally higher MEs than RSPs while smoothing resulted in lower MEs. In the dense arrays (with 100 receivers), ME was consistently lower (Fig. 2C–D). In the dense, random array, the COA algorithm produced the highest MEs (Fig. 2C). RSPs performed better, but was outperformed by all particle algorithm implementations (Fig. 2C). In the dense, regular array, COAs performed more effectively and marginally better than RSPs, but both were outperformed by the particle algorithms (Fig. 2D). In both random and regular dense array designs, particle smoothing lowered ME. Maps from the ACDCPF algorithm exhibited lower ME than those from the ACPF algorithm, but this effect was small. This result is expected in regions with a smooth bathymetry (as simulated here), which effectively spreads out the locational information provided by depth observations. These patterns were borne out by alternative error metrics (Fig. S5) and across all simulated arrays (Figs 2E–F and S6).

**Fig. 2.**
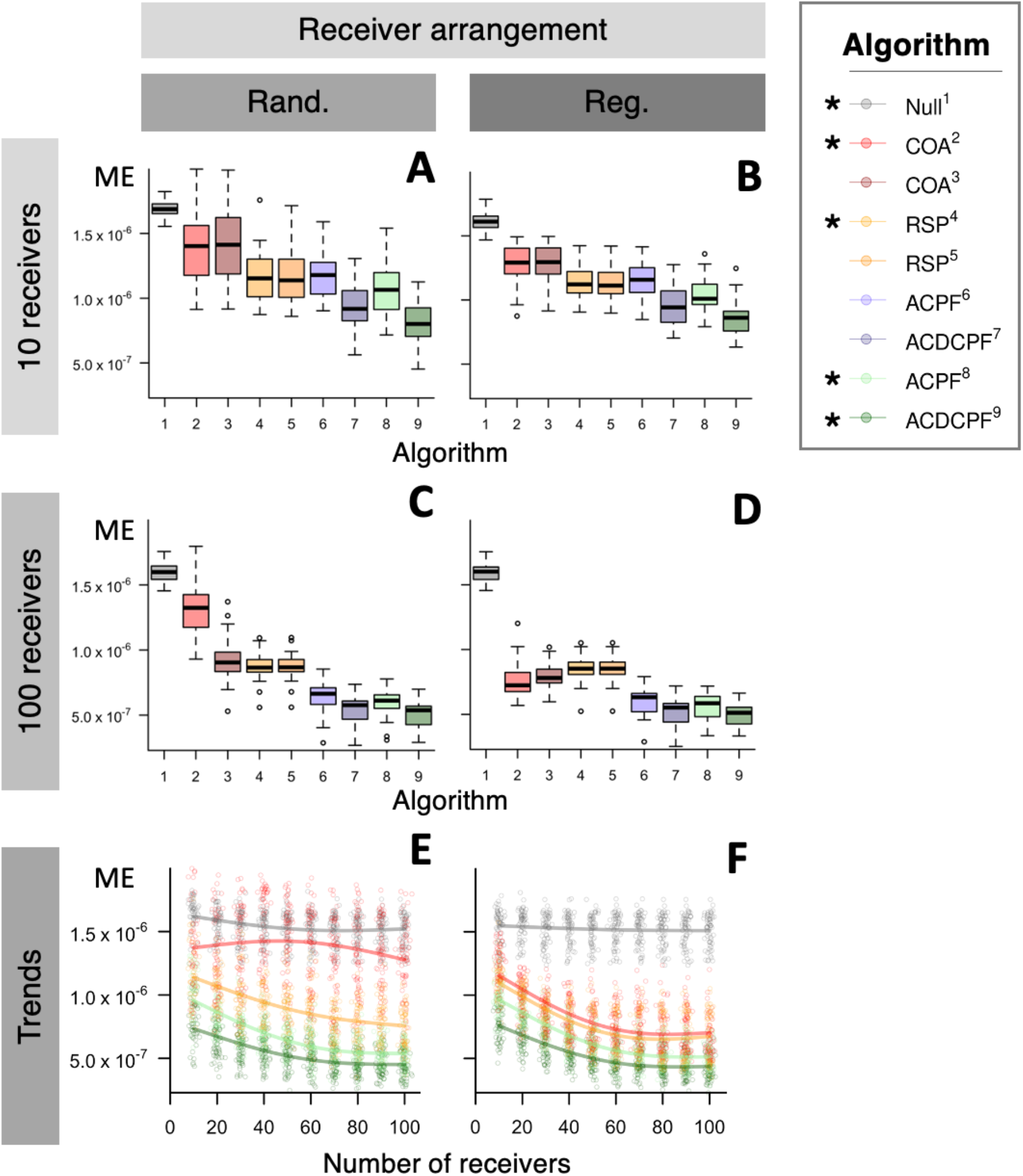
The distribution of mean error (ME) across simulated arrays. Panels **A–D** show the distribution of ME across 30 realisations of the same data generating processes in four example arrays, with 10 or 100 receivers in random or regular arrangements, for nine algorithms. ME was calculated from comparison of each simulated path’s UD (within the duration defined by the first and last detection) and the corresponding UD reconstructed by an algorithm. Boxplot width is proportional to the number of successful algorithm implementations. Algorithm numbers follow Table S4. Panels **E–F** show the trend in ME across array designs with different numbers of receivers for a subset (*) of algorithms. Points mark ME values for specific comparisons and smoothers show trend in the expected ME. All panels correspond to the first simulated study system. For full details, see Figs S5–6.

Out of 1,181 performance simulations, only one ACPF and one ACDCPF simulation failed to converge with the selected number of particles on first implementation. Single-threaded wall times (per time step) for the forward filter averaged 0.011 s for ACPF (5,000 particles) and 0.045 s for ACDCPF (30,000 particles). Wall time for smoothing (1,000 particles) averaged 0.237 s per time step (Fig. S7).

### 3.4. Sensitivity

In the sensitivity analysis, severe parameter mis-specification caused convergence failures (Fig. S8). In the movement process, convergence was most sensitive to under-estimation of mobility, followed by *θ* and *k*, and was insensitive to array design. In the acoustic observation model, convergence failures were associated with overly steep (deflated *α*, inflated *β*), overly shallow (inflated *α*, deflated *β*) and overly truncated (deflated γ) detection probability functions, but the latter two types of mis-specification were most problematic. Overly shallow functions are principally problematic because they excessively restrict particle samples from within detection containers in the gaps between detections (when the influence of *γ* is minimal). In contrast, overly truncated functions are principally problematic at the moment of detection because the *γ* parameter imposes a hard restriction on the region within which particles are sampled (unlike a deflated *α* or inflated *β* value). These effects were generally more common in arrays with more receivers. For both movement and observation parameters, convergence failures were more common for ACDCPF (rather than ACPF) algorithm implementations, with the additional incorporation of depth observations effectively enhancing the chances of prior– data conflicts when the former is mis-specified.

Within the parameter space compatible with the data, parameter mis-specification affected patterns of space use (Figs S9–14). In the movement process, parameter under-estimation produced more concentrated patterns (Figs S9–11). In the observation process, steeper detection probability functions (deflated *α*, inflated *β*) concentrated hotspots around receivers while shallower functions (inflated *α*, deflated *β*) produced depressions in these areas and pitted home ranges (Figs S12–13). (This reflects the way that overly shallow functions restrict particle sampling within detection containers in the gaps between detections, which typically span the majority of a time series.) These effects were more noticeable in regular arrays. Under-estimation of *γ* concentrated patterns around receivers, but over-estimation had a limited effect (Fig. S14). This result fits with the weakly truncating effect of *γ* in our simulations. In all simulations, ACDCPF-derived maps of space use were more robust to parameter mis-specification than maps from the ACPF algorithm (Figs S9–14).

Across the simulations, we observed a U-shaped relationship between ME and the degree of parameter mis-specification, relative to the true value (Figs S15–16). For movement parameters, ME grew more quickly with parameter under-estimation, especially in arrays with more receivers (Fig. S15). Parameter over-estimation, which spreads out the low-probability edges of distributions, produced smaller increases in ME. In the acoustic observation process, ME grew most quickly as detection functions were made shallower (inflated *α*, deflated *β*) and was greatest in arrays with intermediate numbers of receivers (Fig. S16). Under-estimation of *γ* also produced high MEs for the portion of algorithm runs that converged, while over-estimation had little influence.

In almost all cases, mis-specified particle algorithms continued to outperform heuristic methods in terms of ME (Fig. 17).

## 4. Discussion

This study establishes a powerful particle filtering–smoothing methodology for animal movement modelling in autonomous receiver networks such as passive acoustic telemetry systems. The methodology comprehensively represents the movement and detection processes in these systems within a probabilistically sound, flexible and biologically intuitive framework. The process-based perspective is a notable shift from the heuristic methods typically used for movement modelling in passive acoustic telemetry systems (Kraft et al., 2023), which commonly assume direct movements between receivers and analyse properties of the network (rather than the process) (Lédée et al., 2015) or estimate UDs via ‘tuning’ parameters (Niella et al., 2020; Udyawer et al., 2018). The particle methodology reconstructs movements and patterns of space use within and between periods of detection. This produces more accurate maps of space use and facilitates analyses of home ranges, site affinity and habitat preferences. This methodology is a significant development with the potential to support research into the movement ecology and conservation of many species (Hays et al., 2019; Nathan et al., 2022).

The core conceptual advantage of our framework over heuristic approaches is the process-based perspective, which produces outputs with a clear statistical and biological interpretation (Hostetter & Royle, 2020; Lavender et al., 2023). The representation of individual movements is particularly important. Receiver arrays are often irregular and non-overlapping (due to deployment constraints and receiver loss) and ignoring movements or assuming direct transitions between receivers in these settings can suggest overly restrictive patterns of space use that are unduly influenced by array design and for which it is difficult to quantify uncertainty (Lavender et al., 2023). Modelling movements also facilitates analyses of residency—not only around receivers, as quantified by existing indices—but in wider regions of interest (Lavender et al., 2023). This is particularly important where arrays are deployed to inform conservation measures, such as Marine Protected Areas (Lavender et al., 2021; Lea et al., 2016). Representation of the observational processes is also important. In the acoustic telemetry literature, imperfect detectability is acknowledged and quantified using range tests (Kessel et al., 2014), but typically ignored in analyses, which can bias inferences (Winton et al., 2018). Ancillary datasets are also almost exclusively ignored, despite their potential to refine position estimates (Aspillaga et al., 2019; Lavender et al., 2023).

For modelling patterns of space use, our results provide the first substantive assessment of the performance of common heuristic methods and demonstrate the significant benefits of the process-based methodology. For the first time, we clearly demonstrate that the COA algorithm performs poorly relative to alternatives across the board. In sparse, irregular arrays, this algorithm is sometimes worse than a null model and barely improves with receiver number. The widespread adoption of this method and its recent promotion as the centre of a universally applicable analytical framework for passive acoustic telemetry therefore appear misplaced (Udyawer et al., 2018). Particularly in irregular arrays, even a simple representation of movement, as in RSPs, substantially improves maps of space use. For RSPs, this is a reassuring result, given limited evaluation of the method to date. In arrays with more receivers, movement-orientated methods also improve more quickly than the COA algorithm and continue to represent underlying patterns of space use more faithfully, even in dense arrays. These results indicate that, in most real-world settings, the prior (the movement process) contains valuable information and should be represented in analyses. Alongside the COA algorithm, the RSP methodology was consistently outperformed by particle algorithms, especially in irregular arrays. This performance difference was increased by the inclusion of the depth observations, even in the smooth bathymetric landscape we simulated. Given continued improvements in technology and the increasing wealth of data collected alongside acoustic detections (Matley et al., 2023), this is an encouraging result. The integration of state-space models for acoustic detections with ancillary data, achieved here for the first time, holds considerable future promise.

This is not to say that heuristic methods should be superseded by state-space models. Heuristic methods have a track record in the literature (Kraft et al., 2023; Udyawer et al., 2018) and conservation science (Lavender et al., 2021; Lea et al., 2016). They are quick to apply and, as suggested here, may indicate similar patterns of space use to state-space models when data are particularly sparse (and all approaches struggle) or dense (and the prior’s contribution is dampened). In situations with clear expectations, performance can be also improved by tuning parameters on a case-by-case basis. For these reasons, we maintain that different methods are more or less useful in different contexts and caution that analytical standardisation, while valuable in some settings, is not always appropriate (Udyawer et al., 2018). Building on this study, we call for further research into the utility of alternative methods in different settings.

This study unifies and enhances the ‘flapper’ algorithms within a formal particle filtering– smoothing methodology (Lavender et al., 2023). This methodology integrates the so-called AC- and PF-branch algorithms within a single, mathematically coherent framework (Lavender et al., 2023). On a practical level, this reformulation substantially improves algorithm efficiency and facilitates the integration of disparate datasets that vary in quality and resolution. In the flapper algorithms, AC-branch algorithms were required to define the possible locations of an individual, given the data, across a grid at each time step, which becomes prohibitively expensive with increasing grid size. In the particle filter, particle simulation and the likelihood evaluations achieve the same objective but are restricted to particle locations, which removes the dependence on grid size. By recasting the flapper algorithms in their entirety as a particle methodology, we can also start to exploit advanced developments in this field. The substantial refinement of patterns of space use in a coupled filtering—smoothing algorithm demonstrates the potential in this area. This innovation also links movement modelling in acoustic telemetry systems to the handful of particle algorithms developed for animal tracking in other systems and suggests potential refinements (such as two-filter smoothing) that may support applications in those systems (Andersen et al., 2007; Liu et al., 2019).

Our particle methodology is related to existing state-space models in the acoustic telemetry literature (Alós et al., 2016; Hostetter & Royle, 2020; Pedersen & Weng, 2013). The main differences are formulation of the sub-models (which is system-specific), the incorporation of ancillary data (which is theoretically straightforward but only achieved here) and the estimation method (that is, maximum likelihood (Pedersen & Weng, 2013), Markov Chain Monte Carlo (Alós et al., 2016; Hostetter & Royle, 2020) or sequential Monte Carlo (this study)). For example, Alós et al. (2016) and Hostetter & Royale (2020) fit state-space models for acoustic detections using Just Another Gibbs Sampler (JAGS). In theory, this approach benefits from data-driven parameter estimation and directly samples the joint distribution, *f*(***s***_1:*T*_|***y***_1:*T*_). However, JAGS explores correlated distributions inefficiently and can be prohibitively expensive in real-world settings. In contrast, the particle filtering–smoothing methodology targets the simpler distribution *f*(***s***_*t*_|***y***_1:*T*_) but can be orders of magnitude faster (Lavender et al., 2024b).

There are practical challenges to the use of particle filters. The present formulation of our filter requires information on the movement and measurement processes that generate observations. Unlike heuristic methods, where movement capacities and observational processes are enveloped by ‘tuning parameters’ or considered at the interpretation stage, state-space models make the sub-models for these processes are explicit (and context-specific). While available information on these processes can be limited, for the acoustic observation model our simulations show that the shape of the detection probability function is more important than the detection range (in line with the typical sparsity of detections), providing the detection range is large enough. Where data are lacking, we therefore recommend focusing data collection efforts on characterising the shape of the function and setting the detection range to manufacturer specifications. In contrast, for the movement process, an overly flexible movement model (which was associated with lower MEs in our simulations) may be preferable to an overly restrictive one. That being said, the simulations show that even simplified or imperfect representations of the processes that generate observations can substantially refine maps of space use. As more data are collected, such maps can be further refined.

Particle degeneracy is a second challenge that can make convergence hard to achieve with modest (< 1 million) numbers of particles. In this study, convergence issues were negligible, but elsewhere we have experienced challenges coupling sparse acoustic observations with relatively informative archival datasets for benthic species (Lavender et al., in prep). During detection gaps, such situations permit a labyrinth of possible routes that particles have to explore but ultimately render few of them compatible with the data. In these situations, Hamiltonian Monte Carlo (HMC) may be a preferable implementation algorithm (Betancourt, 2017). Like Gibbs sampling, HMC benefits from access to the entire trajectory, but uses derivatives to make proposals, which is more efficient. State-space models for acoustic telemetry have yet to exploit HMC and this is a promising avenue for further research. However, significant theoretical and practical hurdles remain, including the specification of starting values, the incorporation of ancillary datasets, the requirement for smooth likelihood functions, multimodality and the curse of dimensionality. At the current time, estimation of latent locations in complex environments with relatively informative ancillary datasets therefore remains a hard problem for Bayesian sampling methods and alternative approaches may be required (Pedersen et al., 2008). However, in many other situations, including the analyses in this study, these considerations are less relevant because particles can move around available routes more freely (when ancillary data are less informative) or are restricted to fewer possibilities (when ancillary data are highly informative).

A third challenge is that Bayesian techniques can be computationally expensive. The time complexity of the particle filter scales linearly with the number of particles (𝒪(*NT*)) but most smoothing algorithms are more expensive (𝒪(*N*^2^*T*)) (Doucet & Johansen, 2009). In dense receiver arrays where particle trajectories are relatively constrained, our simulations suggest that particle filtering may be sufficient, but in other situations smoothing substantially improves maps of space use. In this study, we averaged 0.01–0.05 s per time step on the particle filter (with 5,000–30,000 particles) and 0.24 s for smoothing (with 1,000 particles) in single-threaded mode on a standard personal computer. While further computational optimisation remains desirable, these speeds compare favourably with related routines for fitting state-space models (Hostetter & Royle, 2020; Liu et al., 2019) and make particle algorithms serious candidates for substantive, real-world analyses (Lavender et al., 2024b).

Looking ahead, we anticipate significant opportunities for continued theoretical and applied work in this area. There is scope to tailor the movement model for different applications through the incorporation of three-dimensional states for demersal and pelagic species (Aspillaga et al., 2019) and behavioural states (Lavender et al., prep). In the representation of the observational processes, one could account for random acoustic transmission intervals, if required (Hostetter & Royle, 2020), and incorporate diverse datasets, such as temperature or salinity (Lavender et al., 2023). The development of multi-resolution models that resolve movements at high resolution in acoustic arrays and use sparse ancillary observations to model larger scale movements, is another important area for future work (Pedersen et al., 2008). Joint inference of movement, observation and state parameters may be also desirable in situations where prior knowledge is limited and data are sufficient. Our particle algorithms stand to benefit from a growing literature, which includes gradient-based methods and other techniques with enhanced convergence properties, as well as novel smoothing approaches (Maken et al., 2022). More broadly, significant work remains to investigate how we can improve, optimise and apply the suite of existing methods in different study systems. We point readers interested in applying our methods to the accompanying software packages (Lavender, 2024a, 2024b; Lavender et al., 2024b).

## Acknowledgments

EL was supported by a postdoctoral researcher position at Eawag, funded by the Department of Systems Analysis, Integrated Assessment and Modelling. We thank Stuart Dennis for scientific computing support.

