## Supporting Information for "Particle algorithms for animal movement modelling in autonomous receiver networks"

### 1. Overview

This paper proposes a state-space model and a particle filtering–smoothing algorithm that reconstructs patterns of space use for tagged animals in passive acoustic telemetry systems. In the Main Text, we develop the model and outline its implementation. In this supporting information, we provide an extended algorithmic explanation of the methodology for mathematical and non-mathematical readers.

### 2. Domain

We consider a simple, two-dimensional, continuous study area (domain). Animal movement is represented as a regular series of discrete steps (from $t = 1$ to $t = T$), with observations ($\boldsymbol{y}$) recorded regularly or irregularly along this timeline[1]. Individual locations[2] are denoted by the two-dimensional vector $\boldsymbol{s}_t = (s_x, s_y)$, where $s_x$ and $s_y$ are continuous. For ease of computation, in some instances, we discretise the domain into $R \times C$ non-overlapping square cells. Each grid cell is denoted by a unique index $I = 1, 2, 3, \dots, R \times C$ and $(s_x^*, s_y^*)$ denotes the coordinates of its geometric centre. A column vector of grid cells is denoted $\boldsymbol{I}$. The functions $\boldsymbol{I} = \text{cell}(\boldsymbol{s}_x, \boldsymbol{s}_y)$ and $(\boldsymbol{s}_x^*, \boldsymbol{s}_y^*) = \text{coord}(\boldsymbol{I})$ provide a mapping between the continuous and gridded domains: $\boldsymbol{I} = \text{cell}(\boldsymbol{s}_x^*, \boldsymbol{s}_y^*)$ defines a column vector of the grid cell/cells ($\boldsymbol{I}$) that contain(s) each coordinate pair $(s_x, s_y)$ in a vector of coordinates $(\boldsymbol{s}_x, \boldsymbol{s}_y)$ and $(\boldsymbol{s}_x^*, \boldsymbol{s}_y^*) = \text{coord}(\boldsymbol{I})$ defines the central coordinates of the grid cell/cells in $\boldsymbol{I}$. We simulate the movement process ($\boldsymbol{s}_t \sim f(\boldsymbol{s}_t \mid \boldsymbol{s}_{t-1})$) in continuous space (see §4.1), but some observational processes ($f(\boldsymbol{y}_t \mid \boldsymbol{s}_t)$) may be defined on the grid (see §4.2). Observations ($\boldsymbol{y}$), contingent upon the individual's location, are recorded at regular or irregular intervals, by receivers and ancillary measuring devices, such as archival (depth) tags. We use $\boldsymbol{y}^{(A)}$ to denote a matrix of acoustic

[1] For example, we may consider a series of two-minute time steps from the start to the end of the study period, along which detections at receivers and/or ancillary observations are recorded at regular or irregular intervals. Detections are typically recorded irregularly but, given an acoustic transmission rate of two minutes or less, we effectively collect acoustic observations (that is, detections or non-detections) at each operational receiver at each time step (alongside ancillary observations). Both detections and non-detections provide information about the location of an individual (see §4.2.2).

[2] For simplicity, we consider individual locations in two-dimensions only, but it is a straightforward extension to model movements in higher dimensions.

observations, with one row for each receiver and one column for each time step, and $y_{k,t}^{(A)} \in \{0, 1\}$ to denote the detection or lack thereof of an individual at receiver $k$ at time $t$. Receiver coordinates are denoted $\boldsymbol{r}_k = (s_x, s_y)$. Depth observations are denoted $\boldsymbol{y}^{(D)}$ and form a row vector with $T$ elements. For a summary of notation, see Table S1.

### 3. Algorithm

Our particle filtering–smoothing algorithm embodies the movement and measurement processes that generate observations and simulates sets of locations ('particles') that are consistent with both processes through time. In this section, we outline the application of the algorithm to passive acoustic telemetry data in non-mathematical language. Full details are provided in sections §4–5.

The forward particle filter moves forwards in time, sampling locations ('particles') at each time step that are consistent with the movement model and the observations up to and including each time point (that is, the filter approximates the partial marginal distribution, $f(\boldsymbol{s}_t \mid \boldsymbol{y}_{1:t})$). We assume that all other parameters are known[3]. Each step comprises four stages:

A. **Simulation.** We simulate a set of candidate locations where the individual could have been located. If the individual's starting location is unknown, at the first time step, particles are simulated from a region within the study area (for instance, around the receiver(s) that recorded the first detection). At subsequent time steps, particles are simulated around previous particles using the movement model. For example, a random walk model can be used to 'kick' previous particles into new locations.

B. **Likelihood.** The probability of the data, given each particle, is evaluated[4]. The probability of acoustic data (detection or non-detection at each operational receiver) depends on array design and detection probability. This is resolved by the equations of the acoustic-container (AC) algorithm, presented by Lavender et al. (2023). The

[3] In typical passive acoustic telemetry settings, it is doubtful that the observations are sufficient to estimate the parameters of the movement and observation models. Here, we therefore assume parameters are known, but analyse algorithm sensitivity to parameter mis-specification in our simulations. Another option is to propagate the uncertainty in parameters by implementing multiple filter runs with different parameter values. It is also possible to infer static parameters in particle filters, but this can be difficult and is not pursued here.

[4] The likelihood measures the statistical agreement between the data and proposal locations. For example, at the moment of a detection, the data (a detection) is generally more probable for acoustic transmissions (that is, for particles) nearer to a receiver. The likelihood quantifies this correspondence and is used to upweight proposals that are more compatible with the data.

probability of ancillary data, such as depth observations, is typically more straightforward and resolved, for instance, by the depth-contour (DC) algorithm (Lavender et al., 2023).

C. **Weights.** Likelihoods are translated into sampling weights that encapsulate the information in the data and the movement model. In general, weights are proportional to likelihoods because the act of kicking particles inherently accounts for the movement process.

D. **Resampling.** Periodically, $N$ particles are re-sampled, with replacement, in line with the weights. The resampling process eliminates particles that are incompatible with available information and increases the frequency of more likely particles.

Smoothing algorithms, such as the two-filter smoother, reweight particles to account all of the data both *up to and beyond* each time step (that is, smoothers approximate the full marginal distribution, $f(\boldsymbol{s}_t \mid \boldsymbol{y}_{1:T})$). For example, the two-filter smoother iteratively updates particle weights, using particles from a forward filter run, a backward filter run and the probability densities of all possible movements between locations (see §5).

### 4. Particle filter

#### 4.1. Simulation

##### 4.1.1. Initialisation

In the particle filter, we approximate the partial marginal distribution $f(\boldsymbol{s}_t \mid \boldsymbol{y}_{1:t})$ using a set of particles[5]. A single particle (location) at time $t$ is denoted $\boldsymbol{s}_{i,t}$. The first step in the particle filter is to simulate initial particles (locations). The animal's initial location may be known or unknown. If the initial location is known, we simply generate $N$ equally weighted copies of

[5] To re-iterate, a 'particle' is simply a location sample. A sample of multiple particles at each time step is generated by the particle filter in such a way as to approximate $f(\boldsymbol{s}_t \mid \boldsymbol{y}_{1:t})$; that is, areas of high probability represented by more particles than areas of low probability at each time step (according to the assumed movement process and the observations up to and including each time step).

that location and proceed to the next time step. Otherwise, we sample $N$ particles from a gridded probability distribution[6], such as:

$$\boldsymbol{s}_{i,t} \sim \text{coord}\big(\text{Uniform}(\{1, 2, \dots, R \times C\})\big). \qquad \text{eqn 1}$$

In practice, it is desirable to restrict initial samples to a sensible area, such as the region defined by acoustic detections[7]. This is achieved by the AC algorithm, which defines containers within with an individual must be located according to receivers (Lavender et al., 2023). If an individual is detected at the start of the time series ($t = 1$), we know that at that moment the individual must be within the maximum detection range of the receiver(s) that recorded the detection(s). Given a detection at receiver $k$, we define a container, $C_A$, within which the individual is located, as a disk, $D$, centred at receiver $k$'s location ($\boldsymbol{r}_k$) of radius $\gamma$ (the detection range), excluding any inhospitable habitats ($U$):

$$C_{A,t=1} = D\big(\boldsymbol{r}_{k,\,t^{(A)}\,=\,1},\ \gamma\big) - U, \qquad \text{eqn 2}$$

where $t^{(A)}$ indexes detections in time[8]. For notational convenience, we assume here that $\gamma$ is a constant. It follows that at the same time step, the individual must be within a finite radius of the receiver(s) ($l$) that recorded the next detection, dependent upon the time between detections and the individual's mobility[9]. This second container is denoted $C_B$ and defined by

$$C_{B,t=1} = D\left(\boldsymbol{r}_{l,t^{(A)}\,=\,2}, \gamma + \Delta\big(t^{(A)} = 1, t^{(A)} = 2\big)\right) - U, \qquad \text{eqn 3}$$

where $\Delta(T_1\,, T_2)$ defines the moveable distance between the time indices $T_1$ and $T_2$ (mobility). At $t = 1$, the individual must be located in the intersection of these two containers[10], i.e.,

$$C_{t=1} = C_{A,t=1} \cap C_{B,t=1}, \qquad \text{eqn 4}$$

which we represent on the grid as $\boldsymbol{I} = \text{cell}(C_{t=1})$. We draw samples from this region accordingly:

$$\boldsymbol{s}_{i,t} \sim \text{coord}\big(\text{Uniform}(\boldsymbol{I})\big). \qquad \text{eqn 5}$$

---

[6] In other words, we sample locations at random (with uniform probability) across the study area. It is convenient to sample starting locations on a grid (such as a bathymetry raster), since likelihood evaluations are often implemented across the same grid (see §4.2).

[7] In mathematical language, we sample initial particles uniformly from the part of the study area with support of the acoustic likelihood.

[8] For notational simplicity only, we imagine that the individual was only detected at a single receiver at any one time in this section.

[9] For example, if the individual can move up to mobility $= 500$ m per time step, and two time steps elapse between the first and second detections, then at the moment of the first detection the individual must be within 1000 m of the detection range (say, $\gamma = 750$ m) of the second receiver.

[10] In other words, at the moment of first detection, if $\gamma = 750$ m, the individual must be 750 m of the first receiver and 1000 + 750 m of the second receiver. The intersection between these two regions contains the possible location(s) of the individual at $t = 1$, according to the AC algorithm, and we sample initial particles from this region.

At the first time step, we then proceed to evaluate the probability of the data given each proposal location (§4.2) and (re)sample $N$ particles (locations) (§4.3–4), which become our starting locations at $t = 2$.

##### 4.1.2. Movement

At subsequent time steps ($t \in \{2, 3, \ldots, T\}$), particles are contingent upon previous locations, in line with the individual's movement capacity. In a standard particle filter, each particle is 'kicked' into a new location[11], which we represent as

$$\boldsymbol{s}_{i,t} \sim f(\boldsymbol{s}_{i,t} \mid \boldsymbol{s}_{i,t-1}). \qquad \text{eqn 6}$$

Under a two-dimensional random walk model, the simulated location $\boldsymbol{s}_{i,t}$ is given by

$$\boldsymbol{s}_t = (s_{x,i,t-1} + d_{i,t} \cos \phi_{i,t}, s_{y,i,t-1} + d_{i,t} \sin \phi_{i,t}), \qquad \text{eqn 7}$$

where $d$ and $\alpha$ are independently distributed random variables that represent step length and turning angle respectively (that is, $d_{i,t} \sim f_d(\boldsymbol{\theta})$ and $\phi_{i,t} \sim f_\phi(\boldsymbol{\theta})$ where $\boldsymbol{\theta}$ generically denotes a parameter vector). Following the example in the Main Text, we could sample step lengths from a truncated gamma distribution, with pre-defined shape ($k$) and scale ($\theta$) parameters and a truncation interval defined between zero and mobility. Typically, $k$ and $\theta$ are set such that movements into nearby locations are more likely than movements further afield, but all movement parameters may be functions of state variables, such as behaviour[12]. For turning angles, a simple choice is to sample angles from a uniform distribution.

In practice, it is sensible to truncate the random walk model by boundary conditions to prevent simulated particles jumping into inhospitable habitats (such as on land). Algorithmically, this is implemented by iteratively simulating $\boldsymbol{s}_{i,t}$ until a valid location is found[13].

[11] What we are doing here is simulating the process of animal movement, in this case using a random walk model in which the animal's location at time $t$ is represented as a function of its previous location, a step length and a turning angle. In the particle filter, we simulate many possible movement trajectories (one for each particle), which are more or less compatible with the data. This is quantified by the likelihood (see §4.2). As the simulation unfolds, we will eliminate trajectories that are incompatible with the data and propagate more likely ones (see §4.2–4).

[12] In the particle filter, the formulation of the movement model is highly customisable. However, particle smoothing requires evaluation of the densities of movements between locations, which places some restrictions on the flexibility of the movement model (see §5).

[13] In the two-dimensional case, for a simulated particle, we simply look up whether or not that particle is valid (i.e., in water). If the particle is invalid, a new particle is simulated. If, even after multiple attempts the simulated location is invalid, we assign that particle zero weight and it is later killed by resampling (see §4.4). See eqns 16–19 for the formal definition of the probability density function from which restricted moves are sampled. (This is required for particle smoothing.)

#### 4.2. Likelihood

##### 4.2.1. Joint likelihood

The next step is to evaluate the probability of the data ($\boldsymbol{y}_t$) given each particle ($\boldsymbol{s}_{i,t}$), i.e.,

$$f(\boldsymbol{y}_t \mid \boldsymbol{s}_{i,t}) = \prod_q f\left(\boldsymbol{y}_t^{(q)} \mid \boldsymbol{s}_{i,t}\right), \quad \text{eqn 8}$$

where $q$ indexes datasets, such as acoustic ($\boldsymbol{y}_t^{(A)}$) and archival ($\boldsymbol{y}_t^{(D)}$) observations[14].

##### 4.2.2. Acoustic observations

###### 4.2.2.1. Contemporary observations

The probability of the acoustic observations depends on detection probability around receivers and array design[15] (Lavender et al., 2023). As explained in the Main Text, we typically assume that detection is a Bernoulli process in which detection probability ($p$) declines logistically with distance between the receiver ($\boldsymbol{r}_k$) and transmitter ($\boldsymbol{s}$). In this instance, the probability of the acoustic observation at receiver $k$ at time $t$ (denoted $y_{k,t}^{(A)} \in \{0,1\}$), given the particle $\boldsymbol{s}_{i,t}$, is defined by

$$\begin{aligned} f\left(y_{k,t}^{(A)} \,\middle|\, \boldsymbol{s}_{i,t}\right) &= \text{Bernoulli}\left(p_{k,t}(\boldsymbol{s}_{i,t})\right) \\ &= p_{k,t}(\boldsymbol{s}_{i,t})^{y_{k,t}^{(A)}} \left(1 - p_{k,t}(\boldsymbol{s}_{i,t})\right)^{1-y_{k,t}^{(A)}} \end{aligned} \quad \text{eqn 9}$$

and

$$p_{k,t}(\boldsymbol{s}_{i,t}) = \begin{cases} \text{logistic}\left(\alpha - \beta \times h(\boldsymbol{s}_{i,t}, \boldsymbol{r}_k)\right) & \text{if } h(\boldsymbol{s}_{i,t}, \boldsymbol{r}_k) < \gamma \\ 0 \text{ otherwise} \end{cases} \quad \text{eqn 10}$$

[14] Particles are more or less compatible with the data and this is quantified by the likelihood. We use this information later to eliminate particles that could not have generated the observations and duplicate particles that could have (see §4.3–4).
[15] Acoustic observations comprise both detections (which are explicitly recorded) and non-detections (which are implicitly known). As a tagged animal moves in an acoustic array, we record detections, typically at irregular intervals. There are many, often unmeasured, variables that determine whether or not detection(s) are recorded (Kessel et al., 2014). For this reason, we treat detection as a stochastic, Bernoulli process in which the probability of a detection at a receiver is specified by a 'detection probability model', such as a truncated logistic distance-decay function (Kessel et al., 2014; Lavender et al., 2023).

where $\text{logistic}(\cdot) = \left(1 + e^{-(\cdot)}\right)^{-1}$; $h(\cdot,\cdot)$ is a distance function between two points, such as the Euclidean distance; α and $\beta$ are (pre-defined) parameters; and $\gamma$ is the assumed detection range[16] (Lavender et al., 2023).

In general, in multi-receiver array with overlapping detection containers, the combined probability of all detection data $f\left(\boldsymbol{y}_t^{(A)} \middle| \boldsymbol{s}_{i,t}\right)$ is given by

$$f(\boldsymbol{y}_t^{(A)} \mid \boldsymbol{s}_{i,t}) = \prod_k f(y_{k,t}^{(A)} \mid \boldsymbol{s}_{i,t}) \qquad \text{eqn 11}$$

where $k$ indexes all operational receivers at time $t$. This follows from the assumption that the detection or non-detection at each receiver is an independent process.

##### 4.2.2.2. Container filter

To facilitate convergence of the filter, it is convenient at this point to eliminate simulated particles that are incompatible with the next detection event. We can use eqn 3 at low computational cost to eliminate invalid particles, since

$$f\left(y_{(k,k+1,\dots,K),\, t^{(A)}+1} \middle| \boldsymbol{s}_{i,t}\right) = \begin{cases} 1 \text{ if } \boldsymbol{s}_{i,t} \in C_{B,t} \\ 0 \text{ otherwise} \end{cases} \qquad \text{eqn 12}$$

where $k, k+1, \dots, K$ index the receivers that recorded detections at the time step of the next detection $(t^{(A)} + 1)$[17].

#### 4.2.3. Ancillary observations

It is straightforward to include ancillary data in the particle filter, with additional likelihood functions. For instance, we can evaluate the probability of depth observations, for both benthic

---

[16] More simply, the likelihood of particles near to receiver $k$ is greater than the likelihood of particles further afield. Particles beyond receiver $k$'s detection range are incompatible with the detection at receiver $k$.

[17] This eliminates proposals that are too far away from the receiver at which the individual was next detected. As in our earlier example (footnote 9, page 4), suppose an individual is detected at $t = 1$ and $t = 3$ and can move up to 500 m per time step. At $t = 1$, any proposals $> 1000 + 750$ m from the next receiver are incompatible with the data: even if the individual was moving at maximum speed, it could not move from those locations into the detection range of the receiver at which it was next detected by the time of the next detection. At $t = 2$, proposals $> 500 + 750$ m from the next receiver are impossible. By $t = 3$, only the particles which have reached the detection container of the next receiver remain valid (from the perspective of the receiver that recorded the detection at $t = 3$). This argument holds for each receiver at which the individual is detected at the next time step, but for notational simplicity we only consider a single such receiver here. In the simulations in this paper, we found that incorporating a container filter, coupled with resampling at every time step, facilitated convergence with substantially fewer particles.

and pelagic species, using the DC algorithm suggested by Lavender et al. (2023) or a similar model. In the Main Text, we present a uniform model that requires $y_t^{(D)}$ to be within an envelope around the bathymetric depth $b(\boldsymbol{s}_i)$ defined by the depth-adjustment functions $\varepsilon_{\text{shallow}}$ and $\varepsilon_{\text{deep}}$, i.e.,

$$f\left(y_t^{(D)} \,\middle|\, \boldsymbol{s}_t\right) = \begin{cases} z_{i,t} & \text{if } b(\boldsymbol{s}_{i,t}) - \varepsilon_{\text{shallow}}(\boldsymbol{s}_{i,t}) \leq y_t^{(D)} \leq b(\boldsymbol{s}_{i,t}) + \varepsilon_{\text{deep}}(\boldsymbol{s}_{i,t}) \\ 0 & \text{otherwise} \end{cases}, \qquad \text{eqn 13}$$

where $z_{i,t}$ is the scaling factor that ensures the distribution normalises to one[18]. This is a slight reformulation of the DC model that permits spatially explicit (e.g., depth-dependent) uncertainty in bathymetric depths. The uncertainty around depth observations is assumed constant and enveloped by the error functions.

### 4.3. Weights

The next step is to define weights with which to (re)sample particles, eliminating unlikely particles and amplifying likely ones. The weight of each particle is given by

$$w_{i,t} \propto w_{i,t-1} f(\boldsymbol{y}_t \mid \boldsymbol{s}_{i,t}) \qquad \text{eqn 14}$$

where $w_{i,t=0} = 1/N$ and $\sum_i w_{i,t} = 1$. This definition holds because we have already accounted for the movement model in the simulation of particles (such that, for instance, particles cluster around previous particles, rather than further afield) and in the resampling step we only need to account for the likelihood.

The effective sample size of particles after weighting is given by

[18] This is an example model for $f(y_t^{(D)} \mid \boldsymbol{s}_t)$ that was motivated by our previous research on flapper skate (*Dipturus intermedius*) (Lavender et al., 2023). In the simulations in this paper, we consider a benthic species like flapper skate and formulate the error functions simply as $\varepsilon_{\text{shallow}}(\boldsymbol{s}_{i,t}) = \varepsilon_{\text{deep}}(\boldsymbol{s}_{i,t}) = 5$. This formulation forces the individual to be in a location where the depth of the seabed ± 5 m overlaps with the observed depth. The constant (5 m) represents the combined accuracy of the observation and the bathymetry data, which are assumed constant (and in our case simulated). In real-world ecosystems, an example formulation of $\varepsilon_{\text{shallow}}(\boldsymbol{s}_{i,t})$ that is more realistic (and similar to that formulated for flapper skate by Lavender and colleagues) is $\varepsilon_{\text{shallow}}(\boldsymbol{s}_{i,t}) = 4.77 + 2.50 + \sqrt{0.5002 + (0.013\, b(\boldsymbol{s}_t)^2} + 5.00$, where 4.77 m is the tag accuracy, 2.50 m is the tidal range, $\sqrt{0.5002 + (0.013\, b(\boldsymbol{s}_t)^2}$ is a depth-dependent bathymetry accuracy (provided by hydrographic surveys) and 5.00 m is an additional term that permits limited movements off the seabed (as hypothesised to occur for flapper skate during prey-strike events). The term $\varepsilon_{\text{deep}}(\boldsymbol{s}_{i,t})$ could be defined similarly (excluding the demersal term). For pelagic species, $\varepsilon_{\text{shallow}}(\boldsymbol{s}_{i,t})$ may further exceed $\varepsilon_{\text{deep}}(\boldsymbol{s}_{i,t})$ and even equal $b(\boldsymbol{s}_{i,t})$ for species that can surface. Such a formulation would permit the individual to be in any location where the depth of the seabed is at least as depth as $b(\boldsymbol{s}_{i,t}) + \varepsilon_{\text{deep}}(\boldsymbol{s}_{i,t})$. Equation 13 is thus a simple but widely applicable model. However, other formulations of $f(y_t^{(D)} \mid \boldsymbol{s}_t)$, including Gaussian or truncated Gaussian formulations, are perfectly possible and may be preferable in some settings.

$$\mathrm{ESS}_t = \left(\sum_i^N \left(w_{i,t}\right)^2\right)^{-1}. \qquad \text{eqn 15}$$

#### 4.4. Resampling

Periodically, $N$ particles are resampled, with replacement and in line with the weights, from the set of simulated particles. Different resampling strategies are available. Multinomial resampling is a simple option in which particles are sampled according to a multinomial distribution of weights. However, systematic resampling has been shown to outperform multinomial resampling and is generally recommended (Doucet & Johansen, 2009). In this paper, we implement systematic resampling when the ESS falls below the number of particles. This amounts to resampling at every time step, which facilitated convergence in our simulations.

### 5. Smoothing

Particle smoothing algorithms use particles from the particle filter (that is, samples from $f(\boldsymbol{s}_t \mid \boldsymbol{y}_{1:t})$) to approximate full marginal distribution, $f(\boldsymbol{s}_t \mid \boldsymbol{y}_{1:T})$. This is achieved by an iterative updating of particle weights such that each weight accounts not only for all of the data up to that time point but all of the future data as well. The two-filter smoother uses particles from a forward run of the particle filter and a backward run (termed the 'backward information filter'). In this algorithm, in words, the smoothed weight of each particle $j$ at time $t$ (that is, $\tilde{w}_{j,t|T}$) is the normalised sum of the product of the probability densities of movement into $j$ from each corresponding particle from the forward filter ($i$) at time $t-1$ (that is, $f(\tilde{\boldsymbol{s}}_{j,t} \mid \boldsymbol{s}_{i,t-1})$), accounting for the weight of that particle ($w_{i,t-1}$) (see Main Text eqns 14–15).

For a restricted random walk (§4.1.2), the probability density of a movement from $\boldsymbol{s}_{i,t-1} \rightarrow \tilde{\boldsymbol{s}}_{j,t}$ is given by

$$f(\tilde{\boldsymbol{s}}_{j,t} \mid \boldsymbol{s}_{i,t-1}) = \begin{cases} \dfrac{1}{z(\boldsymbol{s}_{i,t-1})} f_{\text{unrestricted}}(\tilde{\boldsymbol{s}}_{j,t} \mid \boldsymbol{s}_{i,t-1}) & \text{if } \tilde{\boldsymbol{s}}_{j,t} \text{ is in water} \\ 0 & \text{otherwise,} \end{cases} \qquad \text{eqn 16}$$

where $f_{\text{unrestricted}}(\tilde{\boldsymbol{s}}_{j,t} \mid \boldsymbol{s}_{i,t-1})$ is the probability density of an unrestricted move and $1/z(\boldsymbol{s}_{i,t-1})$ is the normalisation constant. In practice, it is more convenient to specify the density of the step length ($f_d$) and turning angle ($f_\phi$). Using the vector function $\boldsymbol{g}(\tilde{\boldsymbol{s}}_{j,t}, \boldsymbol{s}_{i,t-1}) \rightarrow (d, \alpha)$ as mapping of the jump $\boldsymbol{s}_{i,t-1} \rightarrow \tilde{\boldsymbol{s}}_{j,t}$ expressed in Cartesian coordinates to polar coordinates, where $d$ is the distance of the jump and $\phi$ is the polar angle of the jump. We apply change of variables to determine the density of an unrestricted movement, as follows:

$$f_{\text{unrestricted}}(\tilde{\boldsymbol{s}}_{j,t} \mid \boldsymbol{s}_{i,t-1}) = f_d\left(\boldsymbol{g}(\tilde{\boldsymbol{s}}_{j,t}, \boldsymbol{s}_{i,t-1})\right) f_\phi\left(\boldsymbol{g}(\tilde{\boldsymbol{s}}_{j,t}, \boldsymbol{s}_{i,t-1})\right) \left| \det\left(J_{\boldsymbol{g}(\tilde{\boldsymbol{s}}_{j,t}, \boldsymbol{s}_{i,t-1})}\right) \right|, \qquad \text{eqn 17}$$

where $J_{\boldsymbol{g}}$ is the Jacobian of $\boldsymbol{g}$. For case of the distributions suggested in Main Text eqns 4–5, the density of a simulated step length is given by

$$f_d(d, \phi) = \text{Truncated Gamma}(d; k, \theta, 0, \text{mobility}) \qquad \text{eqn 18}$$

and the density of a simulated angle is given by

$$f_\alpha(d, \phi) = \text{Uniform}(\phi; -\pi, \pi). \qquad \text{eqn 19}$$

Algorithmically, the two-filter smoother proceeds as follows. Starting at the beginning of the time series, for each particle from the backward filter ($\tilde{\boldsymbol{s}}_{j,t}$), we compute the probability densities of movement transitions from each particle from the forward filter at the previous time step ($\boldsymbol{s}_{i,t-1}$), and thus $\widetilde{w}_{j,t|T}$, following Main Text eqns 14–15 and eqns 16–19 above. The normalisation constant $1/z(\boldsymbol{s}_{t-1})$ corresponds to the probability of moving from $\boldsymbol{s}_{t-1}$ into a permissible location (i.e., in water). We compute this constant using a Monte Carlo simulation, assuming a binomial distribution and $\text{Beta}(1, 1)$ prior. The smoothed weights are used to resample $N$ particles, approximating $f(\boldsymbol{s}_t \mid \boldsymbol{y}_{1:T})$. This process eliminates dead ends ('side-branches' of particles that grow during the particle filter that are sooner-or-later rendered incompatible with the data) while mitigating particle degeneracy (by permitting jumps between novel particle pairs).

### 6. Mapping

Particle samples can be used to reconstruct movement paths and patterns of space use. For path reconstruction, a simple option is to join sequential particles (accounting for weights via resampling). This is primarily useful for visualisation, since particles from the two-filter smoother only approximate $f(\boldsymbol{s}_t \mid \boldsymbol{y}_{1:T})$. 'Proper' movement paths are samples from the joint distribution, $f(\boldsymbol{s}_{1:T} \mid \boldsymbol{y}_{1:T})$, which are much more expensive to obtain. One option is to couple multiple forward filters to a backward sampling algorithm, but we leave this possibility for future studies (Doucet & Johansen, 2009).

Our interest in this paper lies in reconstructing emergent patterns of space use (utilisation distributions), for which samples from $f(\boldsymbol{s}_{1:t} \mid \boldsymbol{y}_{1:T})$ are sufficient. As explained in the Main Text, we recommend computing probability-of-use ($P_I$) at particle locations, discretised across a fine-grid, followed by kernel smoothing. We have implemented this approach in the `patter::map_dens()` function, which wraps `spatstat.explore::density.ppp()` (Baddeley & Turner, 2005; Lavender, 2024). Grid resolution is a trade-off between precision and computation time, as kernel smoothing with large numbers of coordinates can be expensive.

For the simulations in this study, we discretised all coordinates (including particles, centres of activity and simulated paths) onto the same 10 x 10 m grid and used the coordinates of the grid cells containing points (and the associated $P_I$ weights) to estimate utilisation distributions by kernel smoothing (see §7). Smoothed kernel intensity functions were estimated by cross validation with edge correction via the `spatstat.explore::density.ppp()` function (Baddeley & Turner, 2005). Intensity (the expected number of points per unit area) was translated into a utilisation distribution (the probability density of points per pixel) for mapping.

### 7. Analyses

In 'performance' analyses, we compared patterns of space use exhibited by simulated paths to those reconstructed by the COA (Simpfendorfer et al., 2002), RSP and particle algorithms (ACPF and ACDCPF). The reconstruction of patterns of space use for simulated paths and each algorithm is explained below.

- **Simulated paths.** For simulated paths, patterns of space use were generated by fitting kernel UDs (KUDs) to path coordinates using cross validation (see §6).
- **COAs.** For the COA algorithm, we estimated COAs using two arbitrary values for the time interval over which detection locations were averaged ($\Delta T \in \{30, 120 \text{ min}\}$) and estimated KUDs from COAs as described above.
- **RSPs.** The RSP methodology was implemented with the simulated detection range ($\gamma$). Two settings were used for the `er.ad` parameter, which tunes the rate at which the

error around interpolated pseudo-relocations increases away from receivers. This parameter does not have a meaningful interpretation, so we arbitrarily used the default value (0.05 x 250 m) and an inflated value (0.10 x 250 m). Other tuning parameters were set to default values. We used the `RSP::dynBBMM()` function to fit UDs to RSPs, as in the original methodology, but resampled (and renormalised) UDs onto our grid.

- **Particle algorithms.** Particle algorithms were implemented using the correct parameter values and 5,000 or 30,000 particles (for ACPF and ACDCPF, respectively). Trials indicated that this was generally sufficient to achieve convergence in these simulations. We implemented systematic resampling at every time step (when ESS was less than the number of particles). Smoothing was implemented using 1,000 particles (which is sufficient to characterise our two-dimensional target distribution, $f((\boldsymbol{s_x}, \boldsymbol{s_y}) \mid \boldsymbol{y}_{1:T})$). In each implementation, we used particles from (a) the forward filter and (b) the two-filter smoother to estimate KUDs. All KUDs were estimated using discretised coordinates for speed.

We compared UDs from the simulated path and each algorithm visually and with standard error metrics, such as the mean error (the mean difference between a 'true' and reconstructed UD, denoted ME). Algorithms were implemented in single-threaded mode on a socket cluster from `R`. Wall time was tracked on a 2023 MacBook Pro (Apple M2 Pro, 32 GB RAM, 12 CPUs).

In 'sensitivity' analyses, we compared the patterns of space use estimated above to those from particle algorithms with mis-specified parameters to analyse algorithm sensitivity (see Main Text §2.3.4 and Table S5).

### 8. Model skill

We quantified the correspondence between simulated paths' UDs and those reconstructed by each algorithm standard error metrics[19] (Lavender et al., 2022). In the following definitions, the terms $O_I$ and $M_I$ denote the probability density in grid cell $I = 1, \ldots, n$ on the 'observed' UD for a simulated path and a 'modelled' UD, respectively. The vectors $\boldsymbol{O}$ and $\boldsymbol{M}$ represent all 'observed' and 'modelled' values.

[19] In performance analyses, all metrics were used. ME emerged as the metric that distinguished between algorithms most effectively. For sensitivity analyses, we therefore focused on this metric only.

A. **Mean Bias (MB).** MB is the mean difference between modelled and 'observed' values, i.e.,

$$\text{MB} = \frac{\sum_{I=1}^{n}(M_I - O_I)}{n} \qquad \text{eqn 20}$$

B. **Mean Error (ME).** ME is the mean absolute difference between modelled and 'observed' values, i.e.,

$$\text{ME} = \frac{\sum_{I=1}^{n}|M_I - O_I|}{n} \qquad \text{eqn 21}$$

C. **Root Mean Square Error (RMSE).** RMSE is the root mean of the squared differences between modelled and 'observed' values, i.e.,

$$\text{RMSE} = \sqrt{\frac{\sum_{I=1}^{n}(M_I - O_I)^2}{n}} \qquad \text{eqn 22}$$

D. **Spearman's rank correlation coefficient (ρ).** This is a measure of the strength and direction of the correlation between modelled and 'observed' values, defined as

$$\rho = \frac{\text{cov}(R(\boldsymbol{O}), R(\boldsymbol{M}))}{\sigma_{R(\boldsymbol{O})}, \sigma_{R(\boldsymbol{M})}} \qquad \text{eqn 23}$$

where is $\text{cov}(\cdot)$ is the covariance function, $R(\cdot)$ ranks the values and $\sigma_{R(\cdot)}$ is the standard deviation of the ranked variable.

E. **Index of Agreement ($\boldsymbol{d}$).** $d$ is a measure of the correspondence between modelled and 'observed' values that varies from zero (no agreement) to one (perfect agreement), defined by Willmott (1982) as

$$d = 1 - \frac{\sum_{I=1}^{n}(M_I - O_I)^2}{\sum_{I=1}^{n}(|M_I - \hat{O}| + |O_I - \hat{O}|)^2}. \qquad \text{eqn 24}$$
