## Supporting Figures for "Particle algorithms for animal movement modelling in autonomous receiver networks"

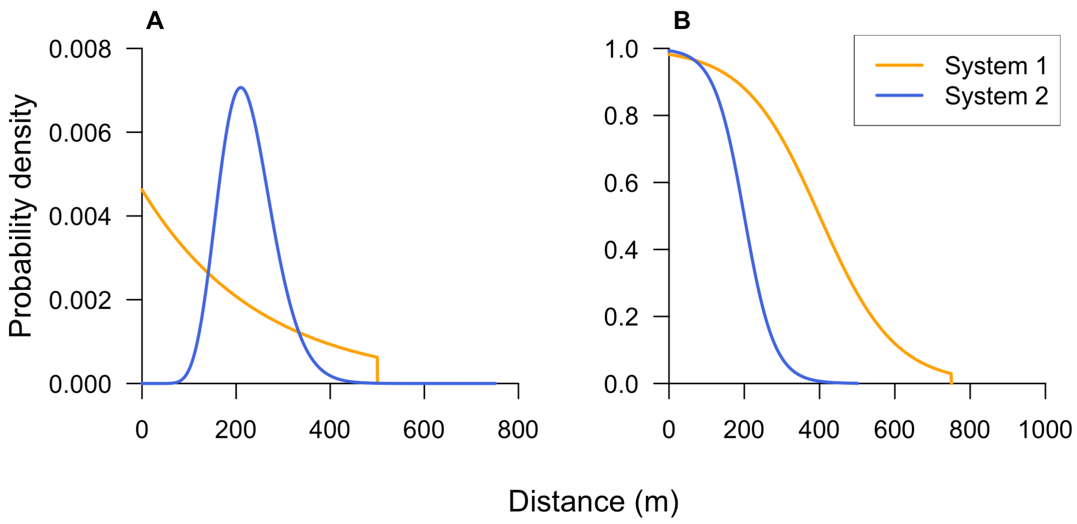


**Fig. S1. True process models for (A) movements and (B) acoustic observations in the two simulated study systems.** **A** shows the probability densities of movement step lengths. **B** shows the probability densities of detections with respect to the Euclidean distance between a receiver and an acoustic transmitter. Turning angles and depth observations were uniformly distributed in all simulations. For parameter values, see Table S2.

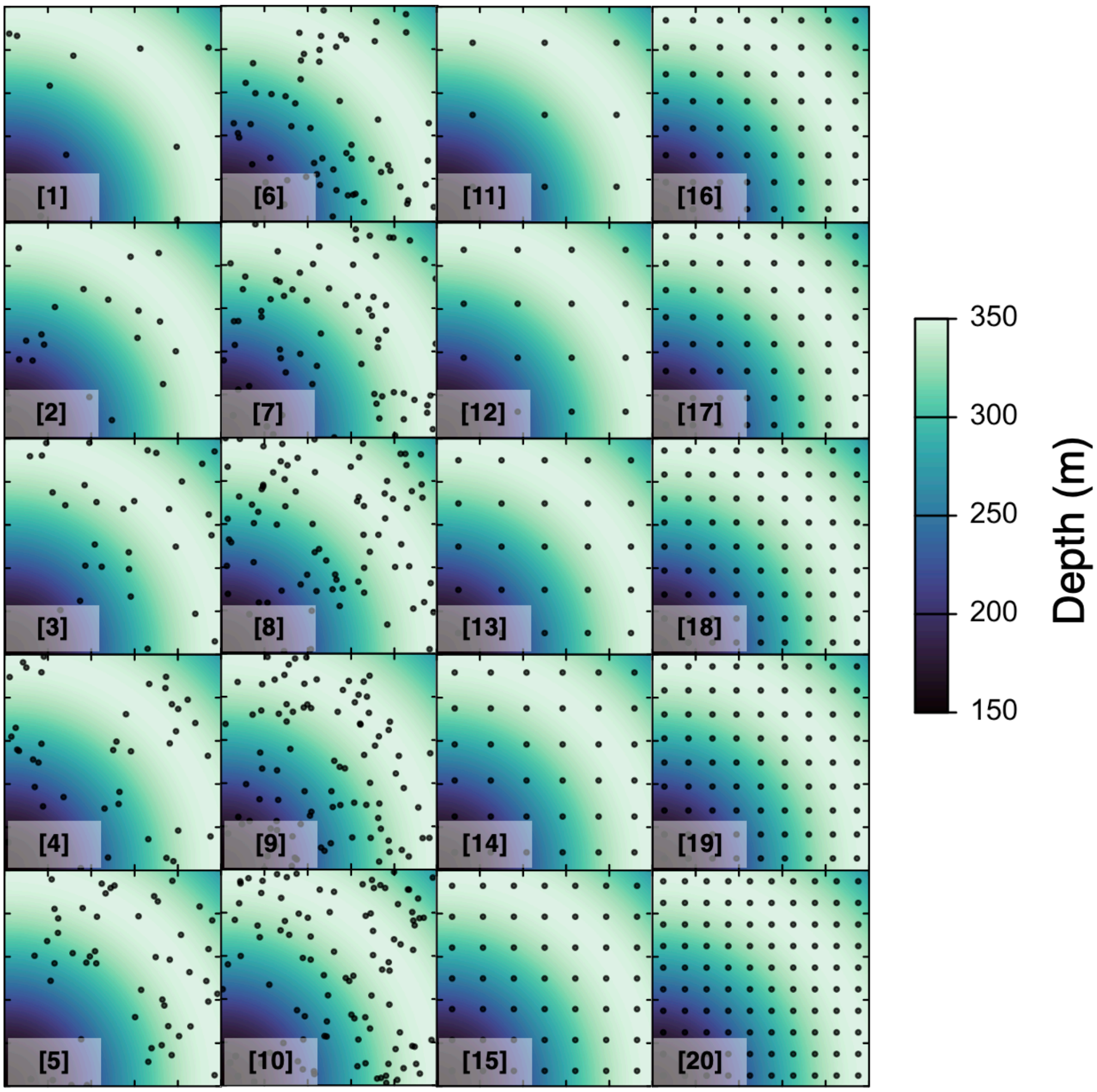


**Fig. S2. Simulated array designs.** Each panel shows a simulated array design. Points mark receivers. Simulated arrays comprised (approximately) $10, 20, \dots, 100$ receivers in random (1–10) or regular (11–20) arrangements (Table S3). The background shows the simulated bathymetry grid (10 x 10 m resolution). The bathymetric depth in each grid cell was defined as $250 - 100 \times \cos\left(\sqrt{((s_x^*)^2 + (s_y^*)^2) / (100\pi)}\right)$, where $s_x^*$ and $s_y^*$ denote the coordinates of the cell's geometric centre (following the notation in Supporting Information §2). Tick marks define 2000 m intervals. The total study area is 100 km$^2$.

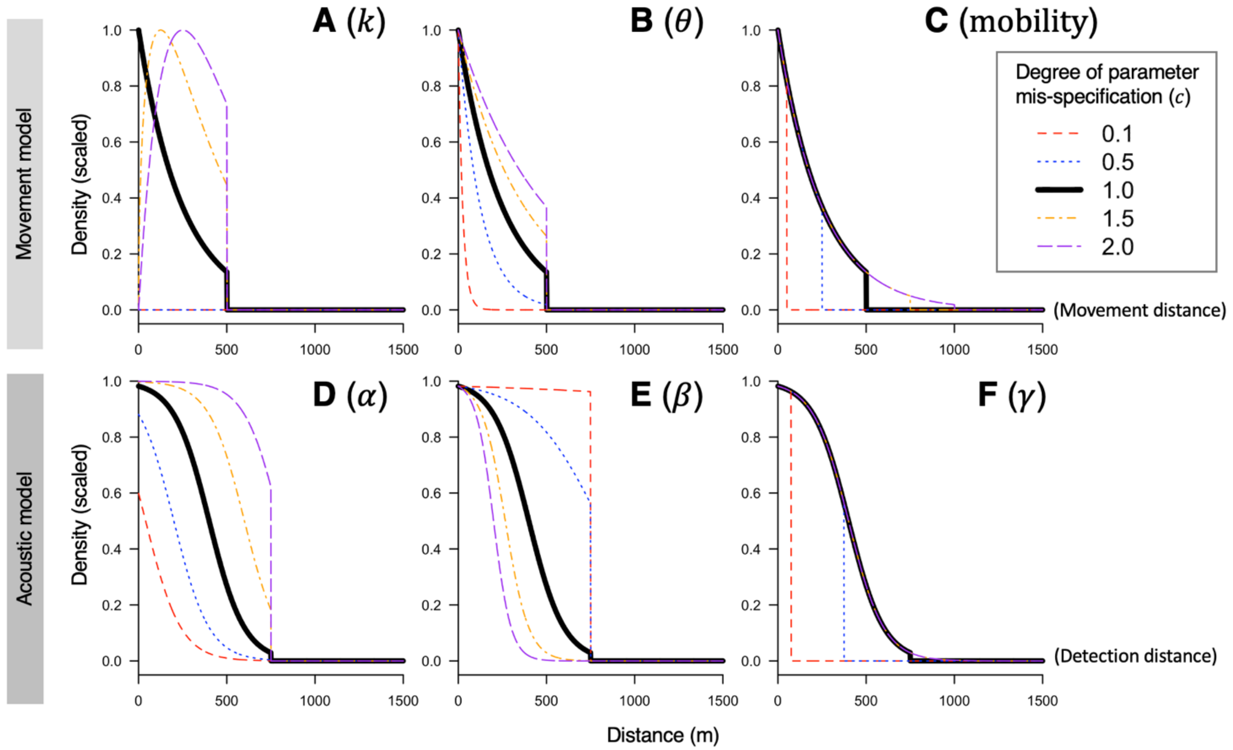


**Fig. S3. Process models for movements (A–C) and acoustic observations (D–F) in sensitivity analyses.** Each panel shows the (scaled) probability density distribution for a selected parameter. Probability densities are scaled to a maximum value of one to facilitate comparisons[1]. For parameter values, see Table S5.

[1] In the first simulated study system, for selected array designs, we re-implemented the particle algorithms using process models with mis-specified parameter values to examine algorithm sensitivity. We examined the influence of mis-specifying selected parameters in the movement model and the acoustic observation model (namely, the $k$, $\theta$ and mobility movement parameters [Main Text eqn 4] and the $\alpha$, $\beta$ and $\gamma$ detection parameters [Main Text eqn 9]). In the truncated Gamma model of step lengths, $k$ and $\theta$ parameters are the shape and scale parameters and mobility is the truncation threshold (i.e., the maximum moveable distance between consecutive time steps). In the truncated logistic detection probability model, $\alpha$ and $\beta$ are the linear coefficients (which determine the shape of the decline in detection probability with distance from a receiver) and $\gamma$ is the truncation threshold (i.e., the maximum detection range, beyond which detection probability is zero).

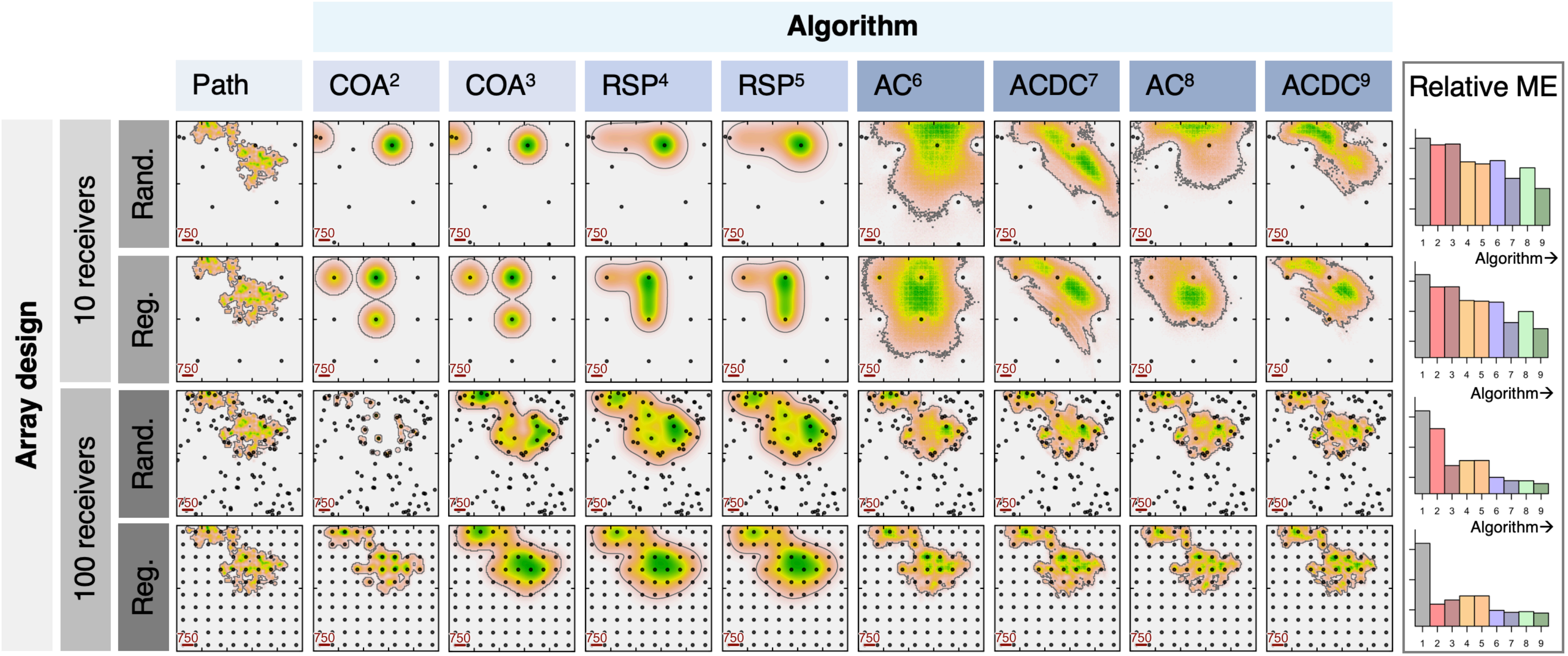


**Fig. S4. Utilisation distributions (UDs) reconstructed by different algorithms in selected array designs for a hypothetical study system.** Following Fig. 1, results are shown for two COA algorithm implementations[2,3], two RSP implementations[4,5], a forward-filter implementation of the ACPF[6] and ACDCPF[7] algorithms and the corresponding filter–smoother implementations[8,9]. For full details, see Table S4. For a simplified version, see Fig. 1.

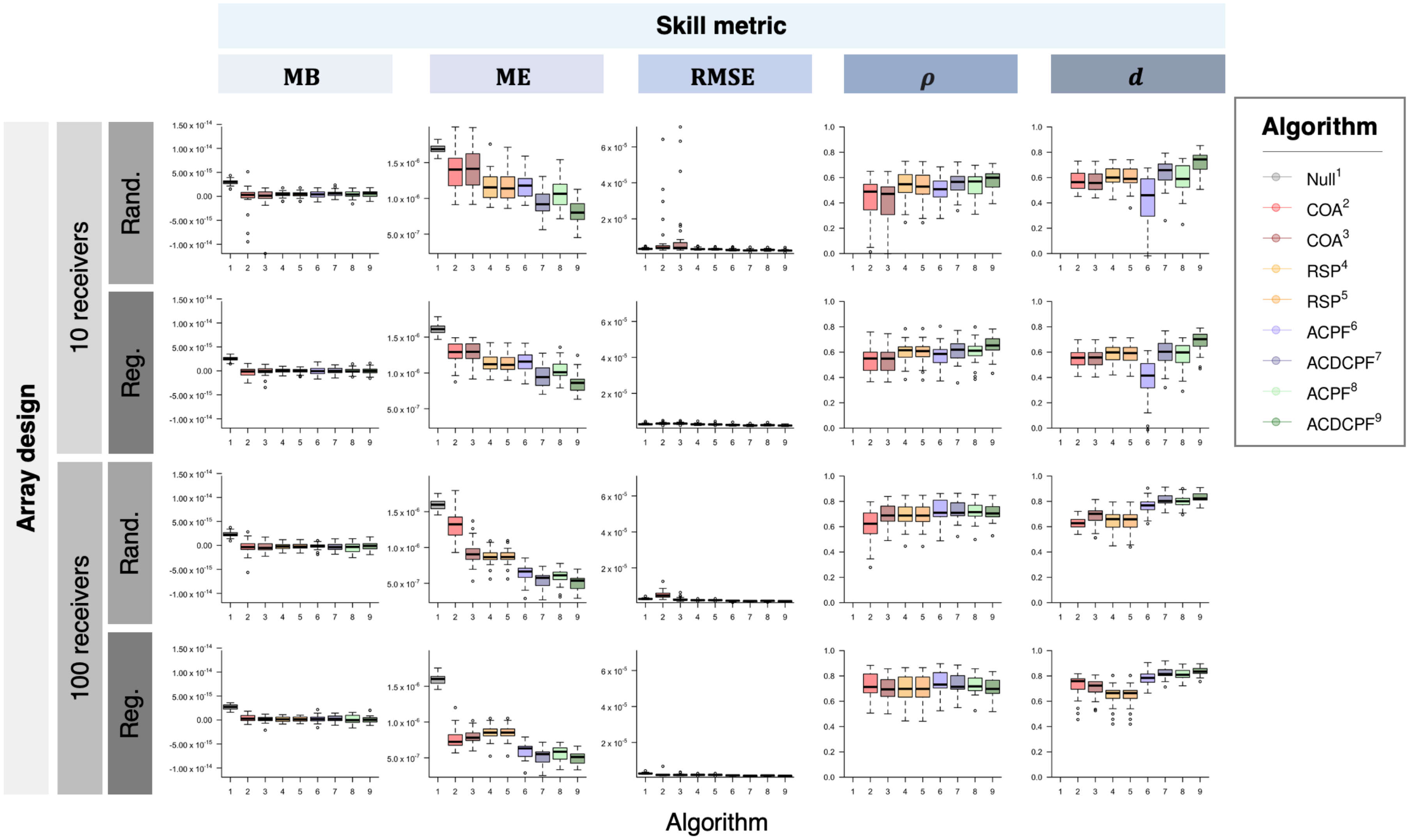

Skill metric
MB
ME
RMSE
ρ
d
Array design
10 receivers
100 receivers
Rand.
Reg.
Rand.
Reg.
Algorithm
Algorithm
Null¹
COA²
COA³
RSP⁴
RSP⁵
ACPF⁶
ACDCPF⁷
ACPF⁸
ACDCPF⁹

**Fig. S5. The distribution of skill metrics in selected arrays for a hypothetical study system.** Four array designs with 10 or 100 receivers in random or regular arrangements are shown. For each array (row), the distribution of skill metrics across 30 realisations of the same data generating processes is shown for two COA algorithm implementations[2,3], two RSP implementations[4,5], a forward-filter implementation of the ACPF[6] and ACDCPF[7] algorithms and the corresponding filter–smoother implementations[8,9] (see Table S4). Skill metrics are derived from comparison of the simulated path's UD (within the duration defined by the first and last detection) and the UD reconstructed by each algorithm. For metric definitions, see Supporting Information §8. $\rho$ and $d$ are undefined for the null model. On boxplots, the thick black line marks the median, the box edges mark the first ($Q_1$) and third ($Q_3$) quartiles and bar ends mark the range (excluding statistical outliers). Points mark statistical outliers (values $< Q_1 - 1.5 \times \text{IQR}$ or $> Q_3 + 1.5 \times \text{IQR}$, where IQR is the interquartile range). Box width is proportional to the number of successful algorithm implementations. Y-axes are constant for each metric but differ between metrics. For a simplified version, see Fig. 2.

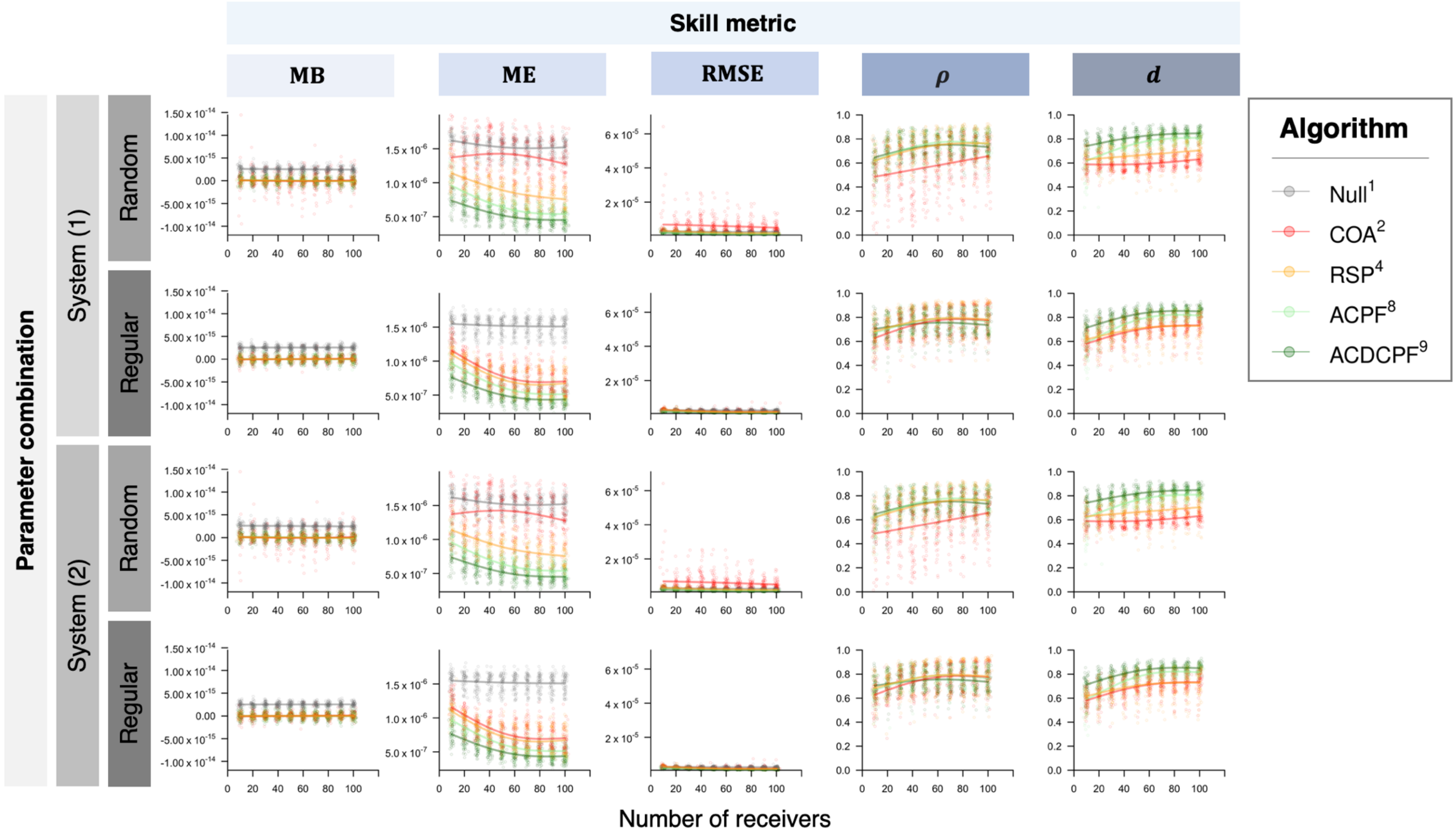
Skill metric
MB
ME
RMSE
ρ
d
Parameter combination
System (1)
System (2)
Random
Regular
Algorithm
Null1
COA2
RSP4
ACPF8
ACDCPF9
Number of receivers

36

**Fig. S6. The distribution of skill metrics for two hypothetical study systems across simulated arrays.** Each panel shows the distribution of skill metrics for different algorithms in a particular hypothetical system, across multiple realisations of the same data generating processes, in random or regularly arranged receiver arrays with different numbers of receivers (Tables S2–4). Metric definitions follow Fig. S5. Points mark metric values for specific comparisons and smoothers show trend in the expected value, estimated for each panel and algorithm by generalised additive models of the form $y_i \sim N(s(a + bx_i), \sigma^2)$, where $\boldsymbol{y}$ represents metric values; $\boldsymbol{x}$ represents the number of receivers; $s$ is a thin plate regression spline with a basis dimension of $k = 3$; $a$, $b$ and $\sigma$ are parameters; and $i$ indexes simulations. Smoothers were generated using `mgcv` (Wood, 2017) and are shown for visual aid only. Consistent results for both study systems shows that patterns are not specific to selected parameter values. For a simplified version, see Fig. 2.

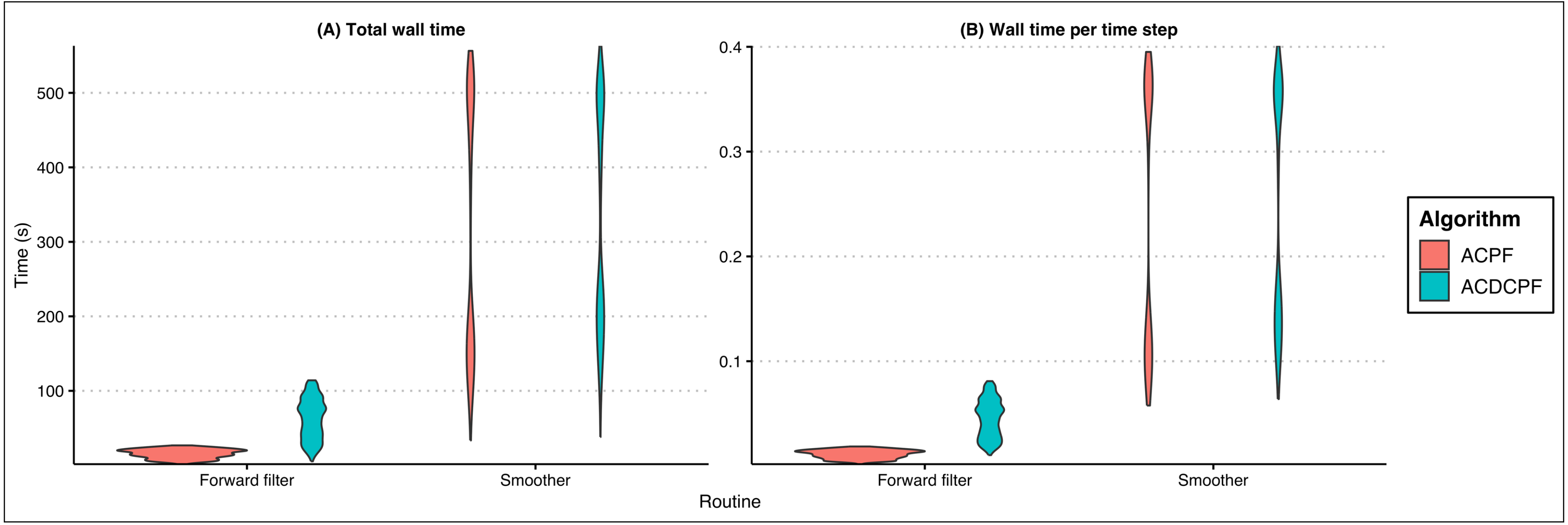


**Fig. S7. Wall time for performance simulations.** Panels (**A**) and (**B**) are violin plots of the total wall time and the wall time per time step for the forward filter and the smoother, respectively. Total wall time is affected by the number of time steps, which differed between simulations (because algorithms were only implemented between the first and last acoustic detections in each simulation). In the forward filter, longer wall times for the ACDCPF algorithm are linked to the larger numbers of particles used (30,000 in ACDCPF versus 5,000 in ACPF) as well as the additional likelihood evaluations required to incorporate depth observations. In both cases, smoothing was implemented with 1,000 particles, so smoothing wall times are similar between the two algorithms. Unlike the forward filter, the time complexity of smoothing is quadratic in the number of particles, and was thus much more expensive in these simulations. Wall time was tracked on a 2023 MacBook Pro (Apple M2 Pro, 32 GB RAM, 12 CPUs). Algorithms were implemented in parallel from `R` in single-threaded mode. Even in single-threaded model, wall times are orders of magnitude faster than comparable routines (e.g., Hostetter & Royle, 2020; Lavender et al., in prep).

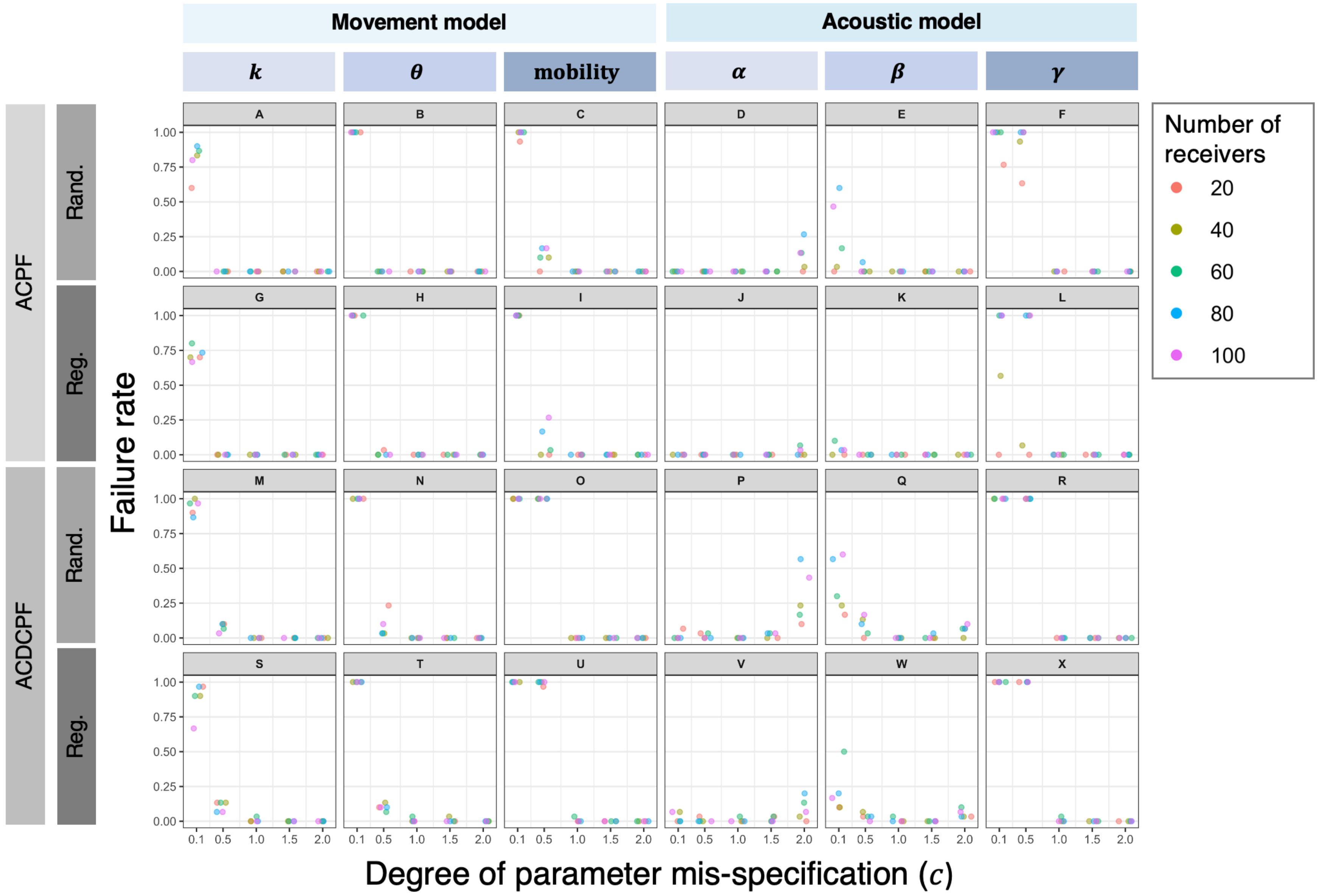

Movement model
Acoustic model
k
θ
mobility
α
β
γ
A
B
C
D
E
F
G
H
I
J
K
L
M
N
O
P
Q
R
S
T
U
V
W
X
ACPF
ACDCPF
Rand.
Reg.
Failure rate
1.00
0.75
0.50
0.25
0.00
0.1
0.5
1.0
1.5
2.0
Degree of parameter mis-specification (c)
Number of receivers
20
40
60
80
100


55

**Fig. S8. Convergence statistics for sensitivity analyses.** Each panel shows the proportion of ACPF or ACDCPF algorithm implementations that failed to converge in sensitivity analyses (simulations with a mis-specified parameter) against the degree of parameter mis-specification for a particular parameter and receiver arrangement (random or regular). For each algorithm and array, proportions were calculated from 30 realisations of the same data generating processes[2]. Points are jittered horizontally for visual clarity. For parameter definitions, see Table S4.

[2] Recall that we simulated 30 realisations of the movement model (i.e., 30 movement paths) and in each of a series of simulated arrays (differing by receiver arrangement and number), we simulated 30 corresponding acoustic and archival datasets. (We did this in two study systems, but given computational restrictions only analysed algorithm sensitivity in the first study system.) In sensitivity analyses, for each simulation (path/dataset), we implemented the ACPF and ACDCPF algorithms with mis-specified parameters. For each simulation, we identified whether or not the algorithm converged and, for the simulations that converged, estimated UDs and ME. This figure shows the proportion of simulations that converged across the 30 simulated paths/datasets in each array (for each algorithm).

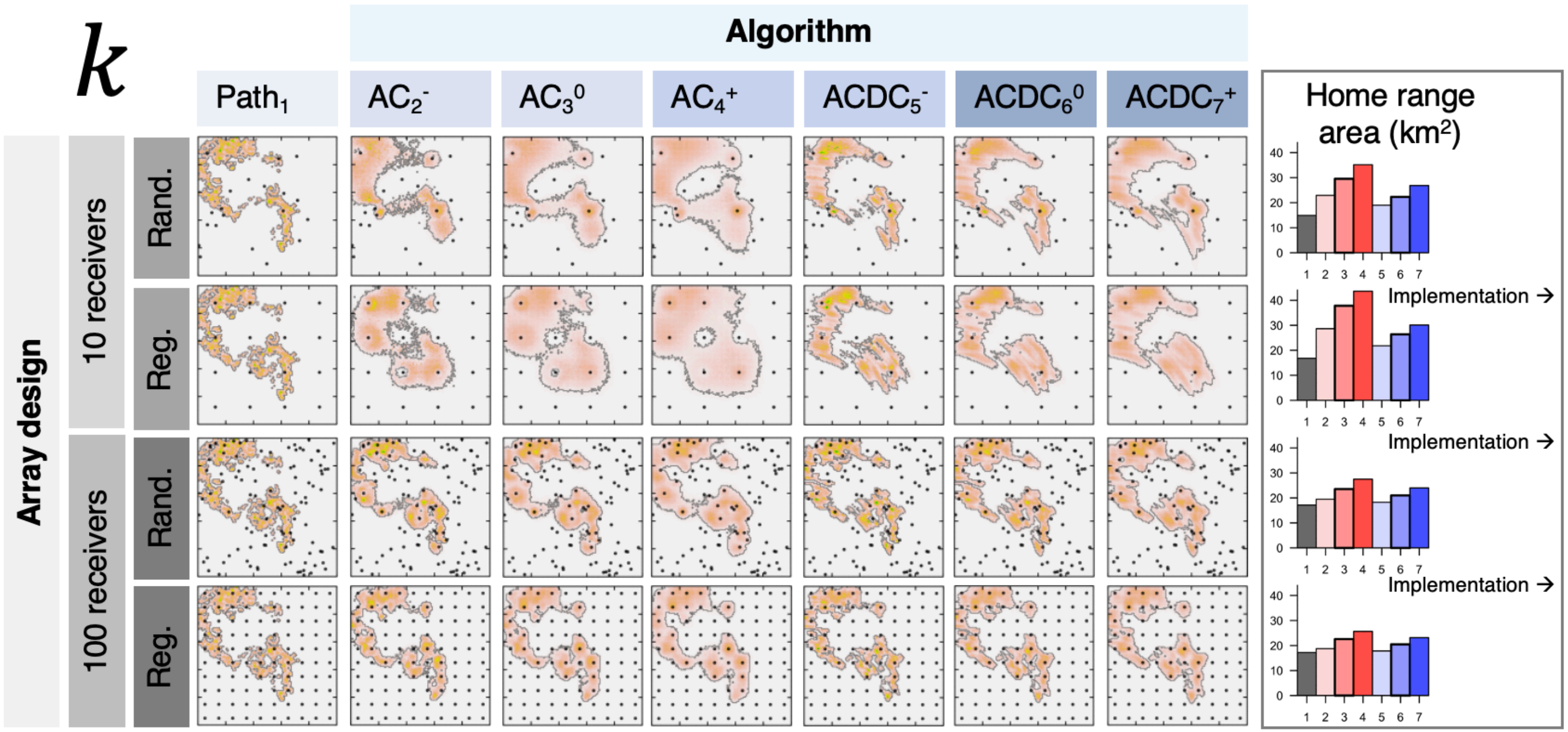


**Fig. S9. Utilisation distributions (UDs) from sensitivity analyses of the movement parameter *k*.** This is the shape parameter of a truncated Gamma distribution of step lengths. Similar to Fig. S4, for the first study system, for selected array designs and path realisations, UDs are shown for the simulated path alongside three ACPF and three ACDCPF two-filter smoother implementations in which the selected parameter was shrunk ($k \times 0.5$, superscript -), held constant ($k \times 1.0$, superscript 0) or inflated ($k \times 1.5$, superscript +) relative the correct value ($k$). For parameter values, see Table S4; for a visualisation of the mis-specified models, see Fig. S3. The map colour scheme follows Fig. S4. Probability densities are scaled to a maximum value of one within rows to facilitate comparisons. Blank map panels denote convergence failures (see Figs S10–14). The area of the home range for each UD is shown by the barplots on the right-hand side. For full details, see Table S5.

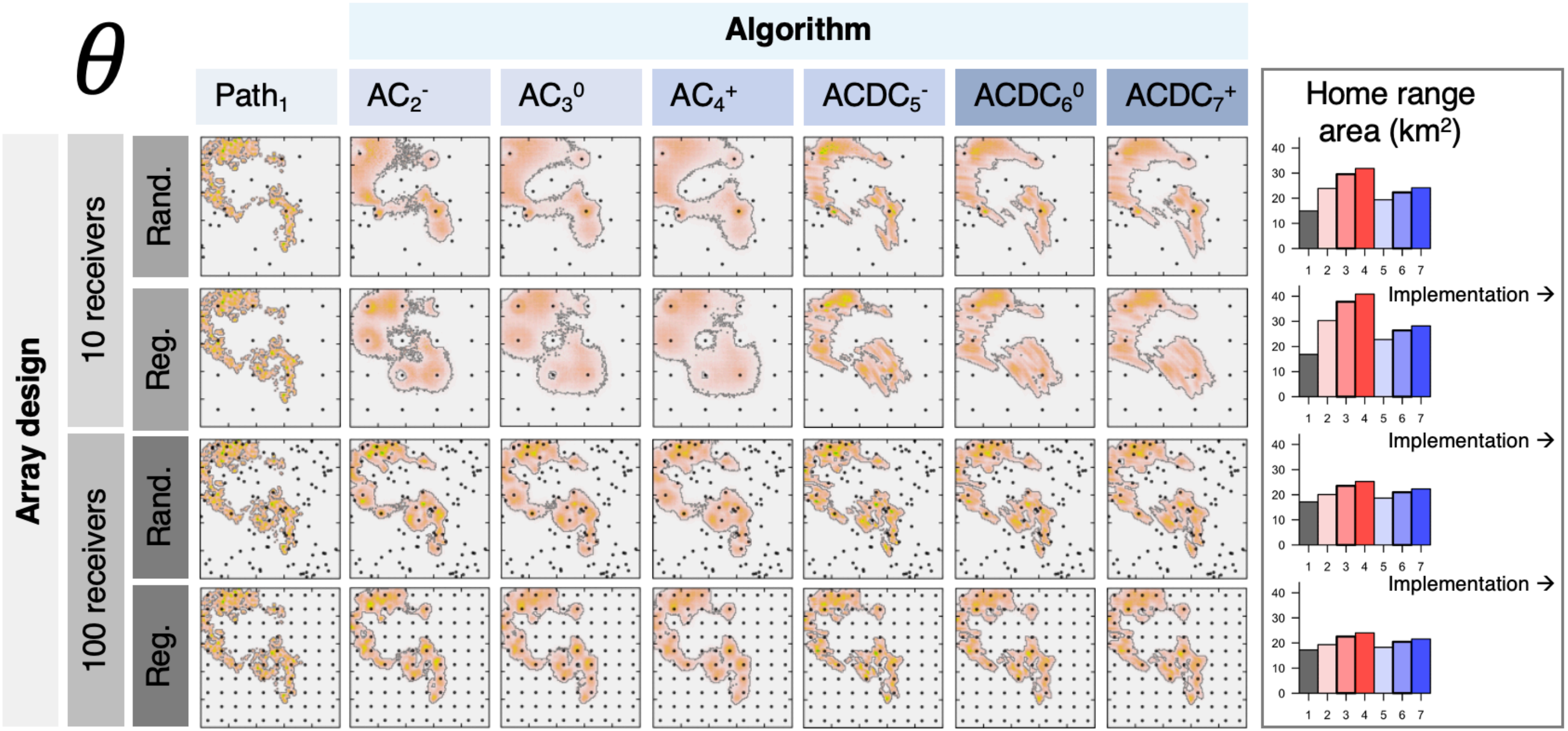


**Fig. S10. Utilisation distributions (UDs) from sensitivity analyses of the movement parameter** $\boldsymbol{\theta}$**, following Fig. S9.** This parameter is the scale parameter of a truncated Gamma distribution of step lengths.

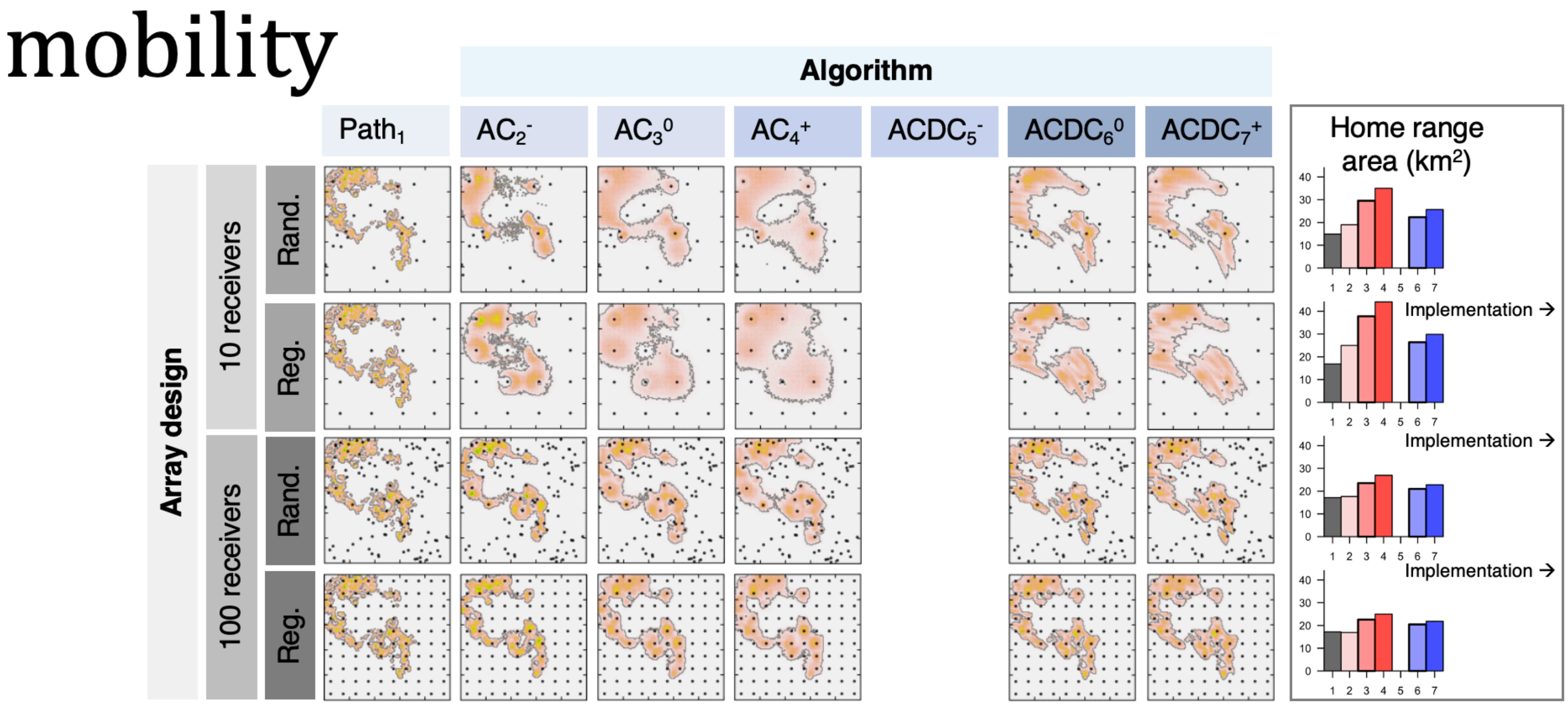


**Fig. S11. Utilisation distributions (UDs) from sensitivity analyses of the movement parameter mobility, following Fig. S9.** This parameter is the truncation parameter in a truncated Gamma distribution of step lengths.

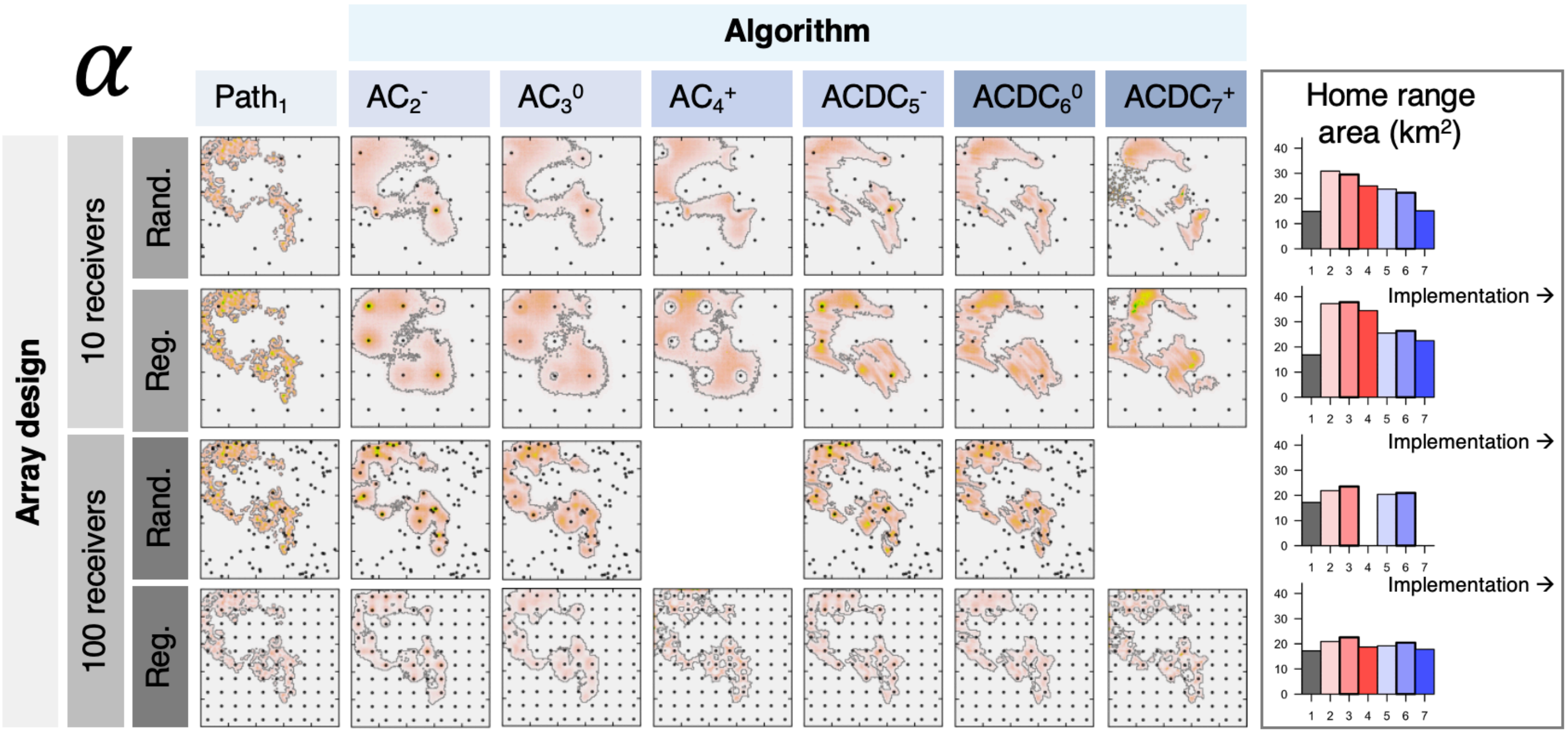


**Fig. S12. Utilisation distributions (UDs) from sensitivity analyses of the acoustic observation process parameter $\alpha$, following Fig. S9.** This parameter is a linear coefficient in a truncated logistic detection probability model. The $\alpha$ and $\beta$ parameters control the shape of the detection probability function with distance from a receiver (see Fig. S3).

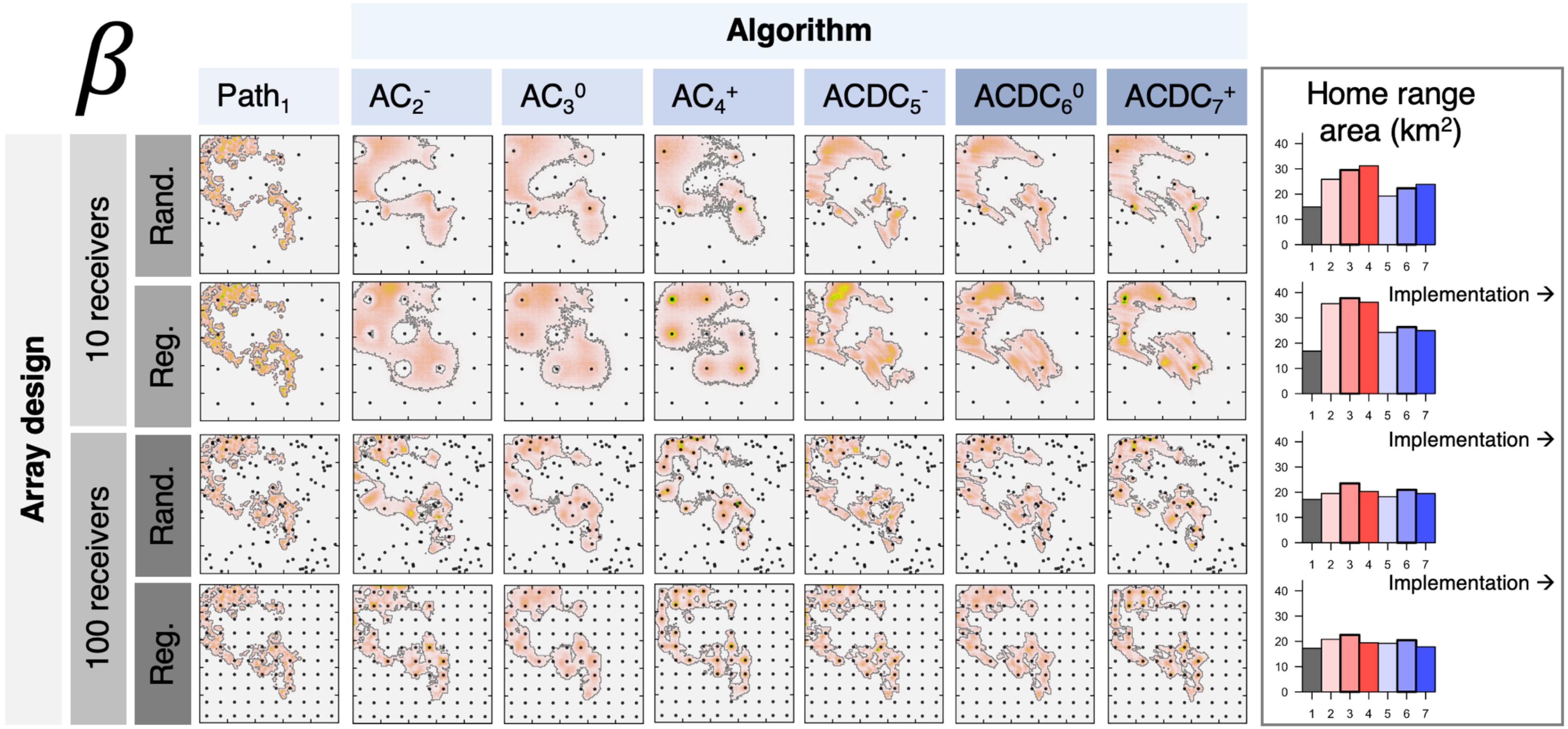


**Fig. S13. Utilisation distributions (UDs) from sensitivity analyses of the acoustic observation process parameter $\beta$, following Fig. S9.** This parameter is a linear coefficient in a truncated logistic detection probability model. The $\alpha$ and $\beta$ parameters control the shape of the detection probability function with distance from a receiver (see Fig. S3).

85

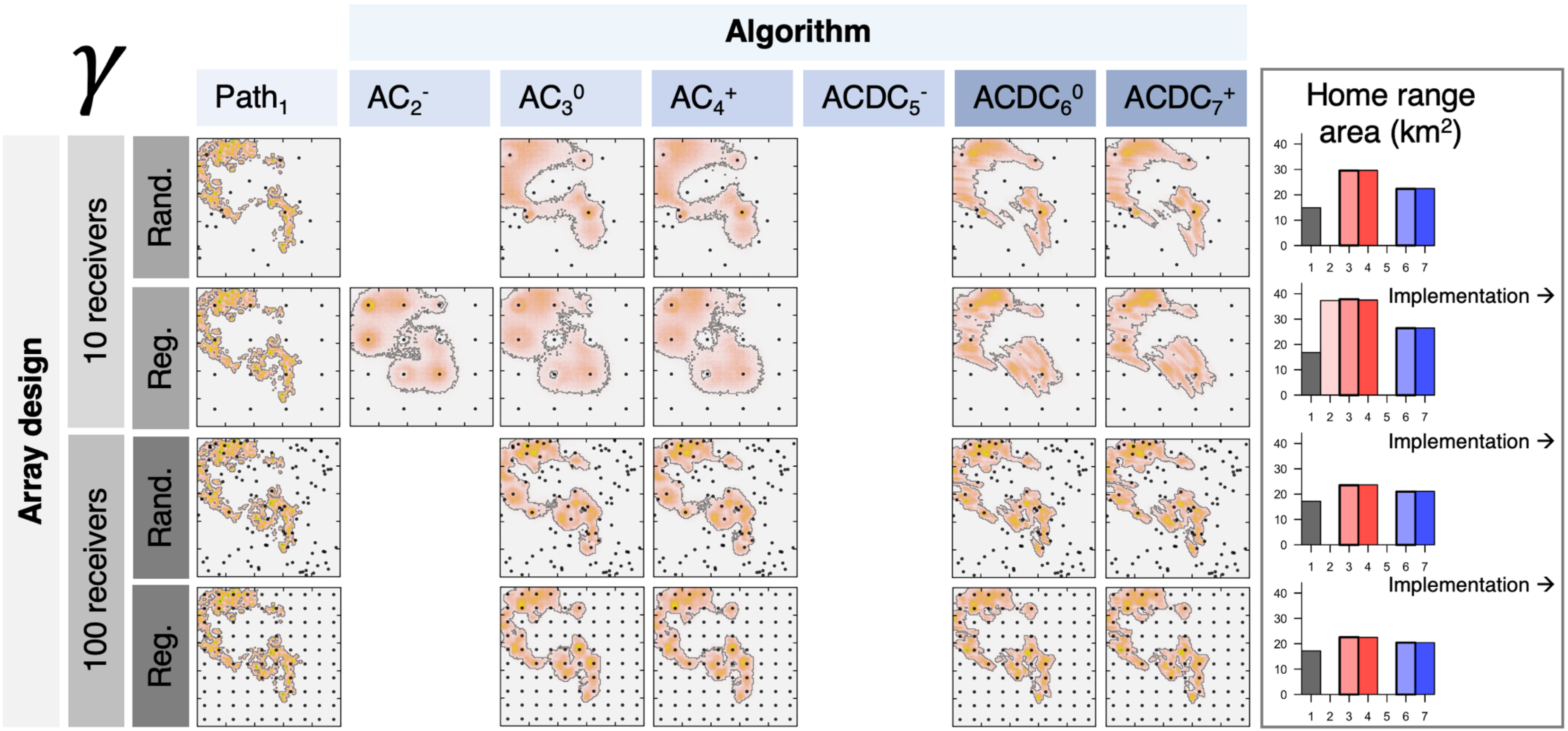


86 **Fig. S14. Utilisation distributions (UDs) from sensitivity analyses of the acoustic observation process parameter $\gamma$, following Fig. S9.** This
87 parameter is the truncation parameter in a truncated logistic detection probability model (i.e., the detection range).

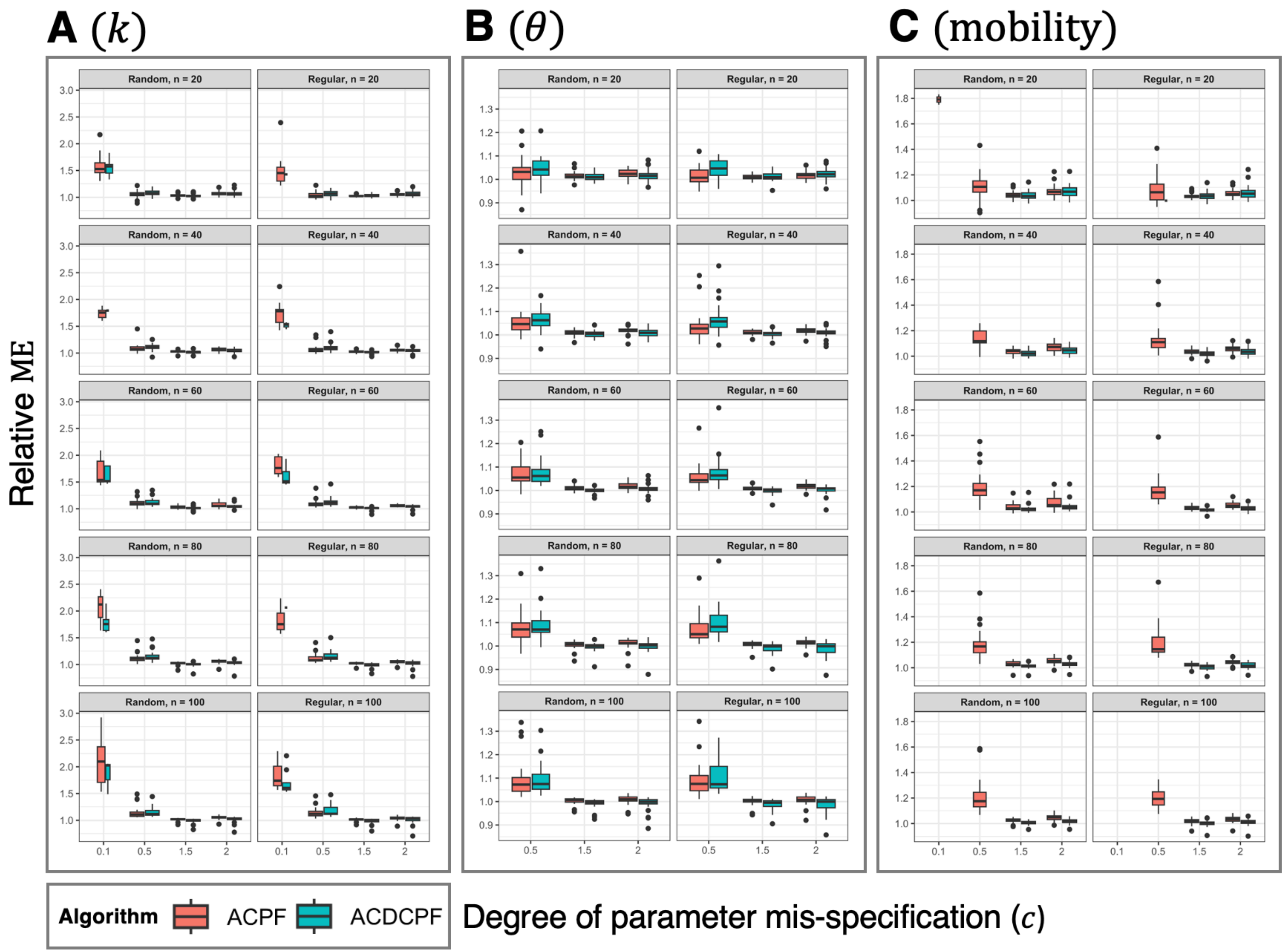
A (k)
B (θ)
C (mobility)
Random, n = 20
Regular, n = 20
Random, n = 40
Regular, n = 40
Random, n = 60
Regular, n = 60
Random, n = 80
Regular, n = 80
Random, n = 100
Regular, n = 100
Relative ME
Algorithm
ACPF
ACDCPF
Degree of parameter mis-specification (c)

**Fig. S15. Algorithm sensitivity to the movement model.** Each box shows the mean error (ME) statistics associated with a particular parameter ($k$, $\theta$ or mobility). Relative ME is the ME of an algorithm implementation (that is, the mean absolute difference between the UD for the simulated path and the reconstructed UD) as a fraction of the ME of the correctly specified algorithm implementation. Within boxes, each panel shows the distribution of relative ME with respect to the degree of parameter mis-specification in a selected array design. Array designs are distinguished by receiver arrangement (random or regular) and number ($n$). For each array, statistics were generated from 30 realisations of the same data generating processes (for further details, see the footnote for Fig. S8). Boxplot properties are as described for Fig. S5. Bar width is proportional to the number of simulations that converged; for some parameters (e.g., mobility), severe parameter under-estimation prevented any simulations from converging and thus calculation of ME. For parameter definitions, see Fig. S3.

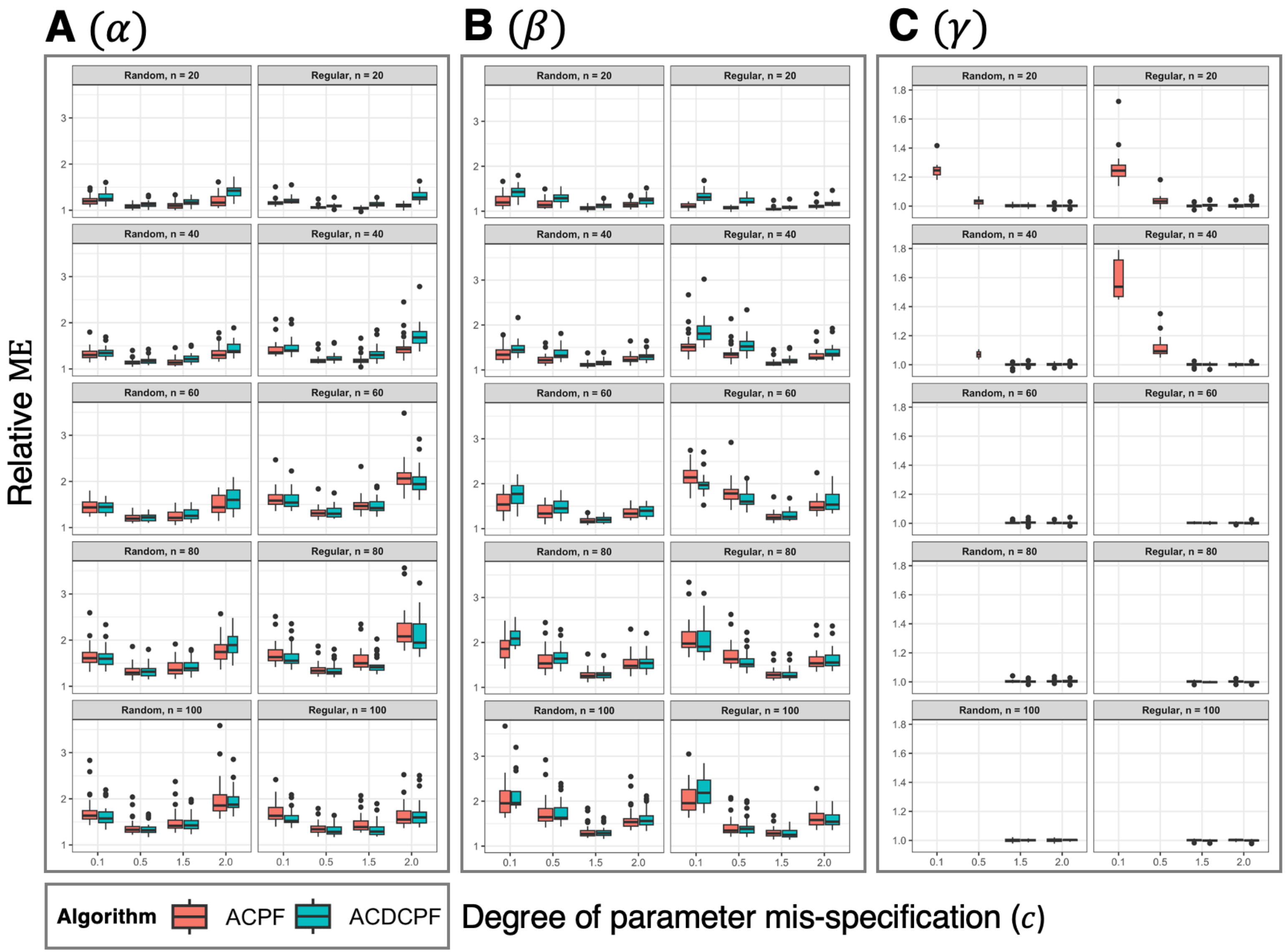


99

100 **Fig. S16. Algorithm sensitivity to the acoustic observation model, following Fig. S15.**

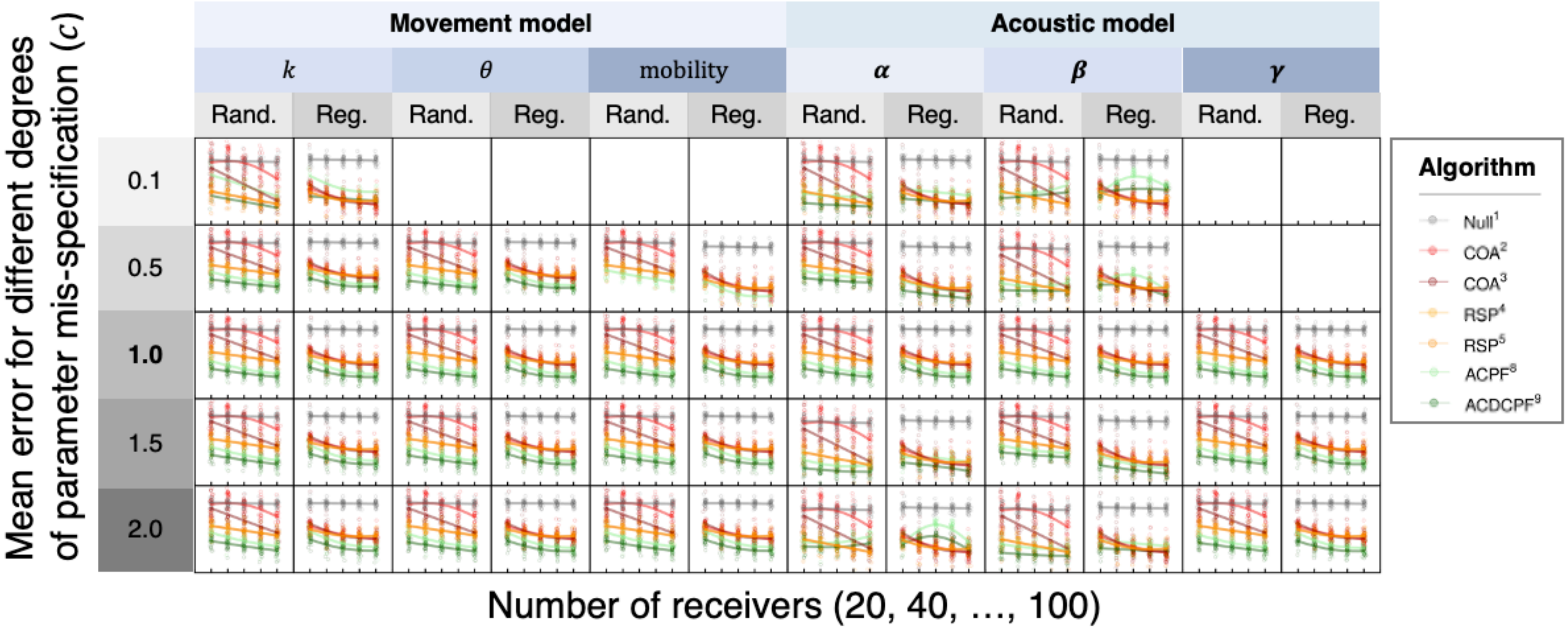
Movement model
Acoustic model
k
θ
mobility
α
β
γ
Rand.
Reg.
0.1
0.5
1.0
1.5
2.0
Mean error for different degrees of parameter mis-specification (c)
Number of receivers (20, 40, …, 100)
Algorithm
Null1
COA2
COA3
RSP4
RSP5
ACPF8
ACDCPF9

101

**Fig. S17. Mean error for heuristic algorithms versus mis-specified particle algorithms.** Similar to Fig. S6, each panel shows, in the first system, for a particular array arrangement (regular or random), the distribution of mean error (y-axis) for different algorithms, across multiple realisations of the same data generating processes, in arrays with different numbers of receivers (x-axis)[3]. Results are shown for a null model, two implementations of the COA and RSP algorithms and an implementation of the ACPF and ACDCPF two-filter smoothing algorithms. For a given receiver arrangement, the data for the null model, COAs and RSPs are the same in all panels. These data are compared against implementations of the ACPF and ACDCPF algorithms with mis-specified parameters. The mis-specified parameter is given by the column and the degree of mis-specification is given by the row. The row $c = 1.0$ corresponds to a correct algorithm implementation (see the panel for ME in Fig. S6). Blank panels correspond to simulations where parameter mis-specification prevented convergence of all ACPF and ACDCPF algorithm runs in a majority of the settings for receiver number (20, 40, …, 100).

[3] Recall that mean error is calculated as the mean absolute difference between the UD for the simulated path and the UD reconstructed by an algorithm. We simulated random and regularly arranged receiver arrays with differing numbers of receivers and in each setting multiple realisations of the same data generating processes. Points mark the mean error for specific comparisons and smoothers show trend in the expected value across all realisations of the same data generating processes (as in Fig. S6). In the figure, it is relative differences between algorithms that are of interest, rather than the absolute values of ME (which are not shown): the figure shows that particle algorithms widely outperform standard heuristic approaches and that this is true even for mis-specified algorithm implementations (except in select, extreme cases of mis-specification). While mis-specified particle algorithms have higher error than correctly specified algorithms (see Figs S15–16), this result suggests that even an imperfect representation of the processes that generate observations can produce maps of space use that outperform those produced by heuristic methods.
