## Supporting Tables for "Particle algorithms for animal movement modelling in autonomous receiver networks"

**Table S1. Summary of notation.** The essential notation in the Main Text and the Supporting Information is listed, ordered alphabetically. Example equation references are provided.

| Symbol | Description |
|---|---|
| $\alpha$ | A linear coefficient in a logistic, distance-decaying, detection-probability model. See Main Text eqn 9. |
| $b(\boldsymbol{s})$ | The bathymetric depth (m) in location $\boldsymbol{s}$. See Main Text eqn 11. |
| $\beta$ | A linear coefficient in a logistic, distance-decaying, detection-probability model. See Main Text eqn 9. |
| $C$ | The number of columns of the grid from which initial particles (locations) are sampled. See Supporting Information eqn 1. The notation $C_t$ is also used to denote acoustic containers but acoustic containers are always subscripted by $t$ (see below). |
| $C_{t=1}$, $C_{A,t=1}$, $C_{B,t=1}$ | Acoustic containers. $C_{t=1}$ defines the set of possible locations of the individual at the initial time step according to acoustic detections (receivers). This is the intersection of the set of possible locations of the individual according to the receiver(s) at which the individual was just detected ($C_{A,t=1}$) and the receiver(s) at which it was next detected ($C_{B,t=1}$). See Supporting Information eqns 2–4. |
| $\text{cell}(\boldsymbol{s}_x^*, \boldsymbol{s}_y^*)$ | A function that maps a vector of coordinate pairs to the vector of indices of cells that contain given coordinates. See Supporting Information eqn 5. |
| $\text{coord}(\boldsymbol{I})$ | A function that maps the vector of cell indices $\boldsymbol{I}$ to a vector of the central coordinates of each cell. See Supporting Information eqn 1. |
| $d$ | The movement step length (m). See Main Text eqn 3. |
| $D(\boldsymbol{r}_{k,\, t^{(A)}=1}, \gamma)$ | A disk around receiver $k$'s location, at the first acoustic time step ($t^{(A)} = 1$), of radius $\gamma$ (the maximum detection range), that defines the individual's possible location according to that receiver (after excluding inhospitable habitats, $U$). See Supporting Information eqn 2. |
| $\delta(\boldsymbol{s}_t - \boldsymbol{s}_{i,t})$ | The Dirac delta function. This is zero when $\boldsymbol{s}_t \neq \boldsymbol{s}_{i,t}$ but integrates to 1 over any region that includes the particle (location) $\boldsymbol{s}_{i,t}$. See Main Text eqn 13. |
| $\delta_{I,I_{i,t}}$ | The Kronecker delta. This evaluates to one if the grid cell of particle $i$ at time $t$ ($I_{i,t}$) equals grid cell $I$ or 0 otherwise. See Main Text eqn 16. |
| $\Delta(T_1, T_2)$ | The maximum moveable distance between two time indices, $T_1$ and $T_2$. See Supporting Information eqn 3. |
| ESS | The effective sample size. See Supporting Information eqn 15. |
| $\varepsilon_{\text{shallow}}(\boldsymbol{s}_t)$ | The shallow-depth adjustment function in the depth-observation model. This returns the maximum possible distance of the individual above the seabed in a given location ($\boldsymbol{s}_t$). See Main Text eqn 11. |
| $\varepsilon_{\text{deep}}(\boldsymbol{s}_t)$ | The deep-depth adjustment function in the depth-observation model. This returns the maximum possible distance of the |

| Symbol | Description |
|---|---|
| | individual below the depth of the seabed in a given location ($\boldsymbol{s}_t$). See Main Text eqn 11. |
| $f(\cdot)$ | A probability density function. See Main Text eqn 1. |
| $f(\boldsymbol{s}_{1:T})$ | The prior probability density (i.e., the movement process), which is modelled as a discrete-time Markovian process. See Main Text eqn 1. |
| $f(\boldsymbol{s}_{t=1})$ | The probability density of the individual's starting location. See Main Text eqn 2. |
| $f(\boldsymbol{s}_t \mid \boldsymbol{s}_{t-1})$ | The probability density of movement from the individual's location at time $t-1$ to the location at time $t$; i.e., the movement model. See Main Text eqn 2. |
| $f(\boldsymbol{s}_t \mid \boldsymbol{y}_{1:t})$ | The partial marginal distribution of the individual's location at a given time ($\boldsymbol{s}_t$) given the data up to and including that time ($\boldsymbol{y}_{1:t}$). This is approximated by the particle filter. See Main Text eqn 12. |
| $f(\boldsymbol{s}_t \mid \boldsymbol{y}_{1:T})$ | The full marginal distribution of the individual's location at a given time ($\boldsymbol{s}_t$) given all of the data ($\boldsymbol{y}_{1:T}$). This is approximated by particle smoothing. See Main Text eqn 15. |
| $f(\boldsymbol{s}_{1:T} \mid \boldsymbol{y}_{1:T})$ | The joint probability distribution of the individual's locations ($\boldsymbol{s}_{1:T}$) and all data ($\boldsymbol{y}_{1:T}$). See Main Text eqn 1. |
| $f(\boldsymbol{y}_t \mid \boldsymbol{s}_t)$ | The probability density of the observations at a given time ($\boldsymbol{y}_t$) given the individual's location at that time ($\boldsymbol{s}_t$); i.e., the likelihood. |
| $f_d(\boldsymbol{\theta})$ | The probability density of step lengths ($d$), where $\boldsymbol{\theta}$ generically denotes a parameter vector. See Supporting Information eqn 18. |
| $f_\phi(\boldsymbol{\theta})$ | The probability density of turning angles ($\phi$), where $\boldsymbol{\theta}$ generically denotes a parameter vector. See Supporting Information eqn 19. |
| $f_{\text{unrestricted}}(\tilde{\boldsymbol{s}}_{j,t} \mid \boldsymbol{s}_{i,t-1})$ | The probability density of an unrestricted movement from location $\boldsymbol{s}_{i,t-1}$ on the forward particle filter to location $\tilde{\boldsymbol{s}}_{j,t}$ on the backward information filter. See Supporting Information eqn 17. |
| $\phi$ | A turning angle (in radians); equivalent of the polar angle. See Main Text eqn 3. |
| $\boldsymbol{g}(\tilde{\boldsymbol{s}}_{j,t}, \boldsymbol{s}_{i,t-1})$ | A vector function $\boldsymbol{g}(\tilde{\boldsymbol{s}}_{j,t}, \boldsymbol{s}_{i,t-1}) \rightarrow (d, \alpha)$ that maps the jump from a location from the forward particle filter to a location on the backward information filter ($\boldsymbol{s}_{i,t-1} \rightarrow \tilde{\boldsymbol{s}}_{j,t}$) expressed in Cartesian coordinates to polar coordinates, where $d$ is the distance of the jump and $\phi$ is the polar angle of the jump. See Supporting Information eqn 17. |
| $\gamma$ | The maximum detection range (m); that is, the distance from a receiver beyond which the detection of an acoustic transmission is assumed impossible. See Main Text eqn 9. |
| $h(\cdot,\cdot)$ | A distance function that evaluates the distance between two locations, such as the Euclidean distance (m). See Main Text eqn 9. |
| $i$ | An index over particles from the forward particle filter. See Main Text eqn 13. |
| $I, \boldsymbol{I}$ | Grid cell indexes. The symbol $I$ indexes over grid cells. See Main Text eqn 16. The symbol $\boldsymbol{I}$ denotes a vector of grid cells. See Supporting Information eqn 5. |
| $j$ | An index over particles from the backward information filter. See Main Text eqn 14. |

| Symbol | Description |
|---|---|
| $J$ | The Jacobian matrix. See Supporting Information eqn 17. |
| $k$ | The shape parameter of a (truncated) Gamma distribution (used to model movement step lengths). See Main Text eqn 4. We also separately use $k$ as an index over operational receivers. |
| mobility | The maximum moveable distance (m) between two consecutive time steps. (This is the upper truncation parameter of the truncated Gamma distribution used to model movement step lengths.) See Main Text eqn 4. |
| $N$ | The total number of particles in the particle filter or the smoother. |
| $p_{k,t}(\boldsymbol{s}_t)$ | The probability of a detection at receiver $k$ at time $t$, given an acoustic transmission from location $\boldsymbol{s}_t$. See Main Text eqn 8. |
| $P_I$ | Probability-of-use for grid cell $I$; that is, the probability that an individual is located in that cell at a randomly chosen time. See Main Text eqn 16. |
| $\boldsymbol{r}_k = (s_x, s_y)$ | Receiver $k$'s two-dimensional location. The expression $\boldsymbol{r}_k = (s_x, s_y)$ denotes that $\boldsymbol{r}_k$ is a two-dimensional vector with x ($s_x$) and y coordinates ($s_y$). See Main Text eqn 9. |
| $R$ | The number of rows of the grid from which initial particles (locations) are sampled. See Supporting Information eqn 1. |
| $\boldsymbol{s}_{i,t}$ | A particle sample at time $t$ from the forward particle filter. See Main Text eqn 13. |
| $\tilde{\boldsymbol{s}}_{j,t}$ | A particle sample at time $t$ from the backward information filter. See Main Text eqn 15. |
| $\boldsymbol{s}_t = (s_x, s_y)$ | The individual's location. The expression $\boldsymbol{s}_t = (s_x, s_y)$ denotes that $\boldsymbol{s}_t$ is a two-dimensional vector with x ($s_x$) and y coordinates ($s_y$). See Main Text eqn 1. |
| $t$ | An index over time steps from $1, 2, \dots, T$. See Main Text eqn 1. |
| $T$ | The final time step. See Main Text eqn 1. |
| $\theta$ | The scale parameter of a (truncated) gamma distribution, used to model movement step lengths. See Main Text eqn 4. |
| $w^*_{i,t}$ | Generic notation for the normalised weight particle $i$ at time $t$ (either $w_{i,t}$, $\widetilde{w}_{j,t}$ or $\widetilde{w}_{j,t\vert T}$). See Main Text eqn 16. |
| $w_{i,t}$ | The normalised weight of particle $i$ at time $t$ from the forward particle filter. See Main Text eqn 13. |
| $\widetilde{w}_{j,t}$ | The normalised weight of particle $i$ at time $t$ from the backward information filter. See Main Text eqn 14. |
| $\widetilde{w}_{j,t\vert T}$ | Smoothing weights in the two-filter smoother. See Main Text eqn 14. |
| $\boldsymbol{y}$ | A dataset of observations. We consider a combined dataset comprising acoustic and archival (depth) observations; i.e., $\boldsymbol{y} = \{\boldsymbol{y}^{(A)}, \boldsymbol{y}^{(D)}\}$. See Main Text eqn 7. |
| $\boldsymbol{y}^{(A)}$ | An $M \times T$ matrix of acoustic observations, which comprise detections (1) and non-detections (0) at each receiver $k$; i.e., $y_{k,t}^{(A)} \in \{0, 1\}$. A column-vector of acoustic observations at time $t$ is denoted $\boldsymbol{y}_t^{(A)}$. See Main Text eqn 7. |

| Symbol | Description |
|---|---|
| $\boldsymbol{y}^{(\boldsymbol{D})}$ | A row vector of depth (m) observations. A single depth observation at time $t$ is denoted $y_t^{(D)}$. See Main Text eqn 7. |
| $\frac{1}{z(\boldsymbol{s}_{i,t-1})}$ | A normalisation constant required to evaluate the probability density of a movement from location $\boldsymbol{s}_{i,t-1} \rightarrow \tilde{\boldsymbol{s}}_{j,t}$. See Supporting Information eqn 16. |

**Table S2. Simulation parameters.** We considered two hypothetical 'study systems' within which we simulated, for a hypothetical individual, movements and observations according to a particular set of movement and observation process parameters[1]. For simplicity, for the movement process, we only considered Truncated Gamma models of step length ($d_t \sim \text{Truncated Gamma}(k, \theta, 0, \text{mobility})$) and a uniform model of turning angle ($\phi_t \sim \text{Uniform}(-\pi, \pi)$). For the acoustic observation (detection) process, we considered a binomial model and a truncated logistic distance-decay model of detection probability, with detection probability declining to zero by $\gamma$ m (Euclidean distance) from receivers. For the archival (depth) observation process, we considered a uniform model: $y_t^{(D)} \sim \text{Uniform}(b(\boldsymbol{s}_t) - 5, b(\boldsymbol{s}_t) + 5)$, where $y_t^{(D)}$ is the depth observation at time $t$ and $b(\boldsymbol{s}_t)$ is the bathymetric depth in location $\boldsymbol{s} = (s_x, s_y)$. In each system, we simulated multiple arrays and realisations of these processes (Table S3). For a visual summary of the movement and detection probability models, see Fig. S1. For 'performance' analyses, simulated datasets were analysed using particle algorithms with the correct parameters (Table S4). For 'sensitivity' analyses, we mis-specified selected parameters, while holding other parameters at the correct values, to examine algorithm sensitivity (Table S5). For parameter definitions, see the Supporting Information §2–5.

| Process | Component | Distribution | Parameter | System | |
|---|---|---|---|---|---|
| | | | | 1 | 2 |
| Movement | Step length ($d$) | Truncated Gamma | $k$ | 1 | 15 |
| | | | $\theta$ | 250 | 15 |
| | | | mobility | 500 | 750 |
| | Turning angle ($\phi$) | Uniform | $a_\phi$ | $-\pi$ | |
| | | | $b_\phi$ | $\pi$ | |
| Acoustic | Detection record $\left(y_{k,t}^{(A)}\right)$ | Binomial | $\alpha$ | 4 | 5 |
| | | | $\beta$ | -0.010 | -0.025 |
| | | | $\gamma$ | 750 | 500 |
| Archival | Depth measurement $\left(y_t^{(D)}\right)$ | Uniform | $a_{y^{(D)}}$ | $b(\boldsymbol{s}_i) - 5$ | |
| | | | $b_{y^{(D)}}$ | $b(\boldsymbol{s}_i) + 5$ | |

[1] We repeated 'performance' analyses for both systems to validate the robustness of resultant patterns to the specific simulation parameters we selected.

**Table S3. Simulated array designs.** For each array, the identifier, receiver arrangement, simulated number of receivers[2] and coverage, for each selected detection range ($\gamma$) parameter, is shown (see Table S2). Coverage is the area contained by (circular) receiver detection containers, expressed as a percentage of the total study area. All arrays were included in 'performance' (P) analyses (see also Table S4). Given computational restrictions, only every other array was included in 'sensitivity' (S) analyses (see also Table S5). For a visualisation, see Fig. S2.

| Identifier | Arrangement | Receiver | Coverage (%): $\gamma$ = 750 m | Coverage (%): $\gamma$ = 500 m | Analyses |
|---|---|---|---|---|---|
| 1 | Random | 10 | 14.68 | 6.88 | P |
| 2 | Random | 20 | 29.06 | 14.38 | P, S |
| 3 | Random | 30 | 40.96 | 21.34 | P |
| 4 | Random | 40 | 43.91 | 24.82 | P, S |
| 5 | Random | 50 | 54.15 | 31.27 | P |
| 6 | Random | 60 | 60.20 | 36.34 | P, S |
| 7 | Random | 70 | 68.77 | 41.76 | P |
| 8 | Random | 80 | 73.46 | 46.45 | P, S |
| 9 | Random | 90 | 75.03 | 49.01 | P |
| 10 | Random | 100 | 79.79 | 53.30 | P, S |
| 11 | Regular | 10 | 15.90 | 7.06 | P |
| 12 | Regular | 20 | 28.26 | 12.55 | P, S |
| 13 | Regular | 30 | 44.16 | 19.61 | P |
| 14 | Regular | 40 | 63.59 | 28.24 | P, S |
| 15 | Regular | 50 | 84.45 | 38.44 | P |
| 16 | Regular | 60 | 95.11 | 50.21 | P, S |
| 17 | Regular | 70 | 95.11 | 50.21 | P |
| 18 | Regular | 80 | 99.59 | 63.54 | P, S |
| 19 | Regular | 90 | 99.59 | 63.54 | P |
| 20 | Regular | 100 | 100.00 | 78.43 | P, S |

[2] The simulated number of receivers is only approximately equal to the realised number of receivers in regular arrays due to the constraints imposed by regularity.

**Table S4. Algorithm implementations for 'performance' analyses.** For each hypothetical study system (Table S2), for selected arrays (Table S3), for each realisation of the data generating processes, we compared patterns of space use for simulated paths to those reconstructed by nine algorithms, including a null model (flat UD); two implementations of the COA algorithm; two implementations of the RSP algorithm; and the forward particle filter and filter–smoother implementations of the particle algorithms (AC* and ACDC*). Parameters for the COA and RSP algorithms, which are heuristic methods, are necessarily arbitrary (apart from $\gamma$ in the case of the RSP algorithm). Correct parameters were used for particle algorithm implementations in performance analyses (Table S2).

| Algorithm | ID | Parameters |
| --- | --- | --- |
| NULL | 1 | - |
| COA | 2 | • $\Delta T = 30$ min |
| | 3 | • $\Delta T = 120$ min |
| RSP | 4 | • $\text{time.step} = 10$ (default)<br>• $\text{distance} = 250$ (default)<br>• $\text{min.time} = 10$ (default)<br>• $\text{max.time} = 48$<br>• $\text{er.ad} = 0.05 \times 250$<br>• $\gamma$ (Table S2) |
| | 5 | • $\text{er.ad} = 0.10 \times 250$<br>• other parameters as above |
| ACPF (forward filter) | 6 | • Table S2 |
| ACDCPF (forward filter) | 7 | |
| ACPF (filter–smoother) | 8 | |
| ACDCPF (filter–smoother) | 9 | |

**Table S5. Algorithm implementations for 'sensitivity' analyses.** For the first hypothetical study system (Table S2), for selected arrays (Table S3), for each realisation of the data generating processes, we compared patterns of space use for simulated paths to those reconstructed by multiple implementations of the ACPF and ACDCPF algorithms with both correct parameter values (Table S4) and progressively mis-specified parameters (to analyse algorithm sensitivity). We mis-specified selected parameters ($p \in \{k, \theta, \text{mobility}, \alpha, \beta, \gamma\}$) via $p^* = cp$, where $c \in \{0.1, 0.5, 1.0, 1.5, 2.0)$, while holding other parameters at the correct values[3] (Table S2). The parameter $c$ is termed the 'degree of parameter mis-specification'; $c < 1$ reflects parameter under-estimation, $c = 1$ leaves parameters unchanged from the correct values (highlighted in grey in the table) and $c > 1$ reflects parameter over-estimation. For a visualisation, see Fig. S3.

| Process | Component | Distribution | Parameter | Degree ($c$) | Value |
|---|---|---|---|---|---|
| **Movement** | Step length ($d$) | Truncated Gamma | $k$ | 0.1 | 0.1 |
| | | | | 0.5 | 0.5 |
| | | | | 1.0 | 1.0 |
| | | | | 1.5 | 1.5 |
| | | | | 2.0 | 2.0 |
| | | | $\theta$ | 0.1 | 25 |
| | | | | 0.5 | 125 |
| | | | | 1.0 | 250 |
| | | | | 1.5 | 375 |
| | | | | 2.0 | 500 |
| | | | mobility | 0.1 | 50 |
| | | | | 0.5 | 250 |
| | | | | 1.0 | 500 |
| | | | | 1.5 | 750 |
| | | | | 2.0 | 1000 |
| | Turning angle ($\phi$) | Uniform | $a_\phi$ | 1.0 | $-\pi$ |
| | | | $b_\phi$ | 1.0 | $\pi$ |
| **Acoustic** | Detection record $\left(y_{k,t}^{(A)}\right)$ | Binomial | $\alpha$ | 0.1 | 0.4 |
| | | | | 0.5 | 2.0 |
| | | | | 1.0 | 4.0 |
| | | | | 1.5 | 6.0 |
| | | | | 2.0 | 8.0 |
| | | | $\beta$ | 0.1 | -0.001 |
| | | | | 0.5 | -0.005 |

[3] Given computational limitations, we focused sensitivity analyses on (a) the first study system, (b) a subset of arrays and (c) selected parameters. In each simulation, we mis-specified a single parameter while holding other parameters constant at the correct values. We did not evaluate the consequence of simultaneous mis-specification of multiple parameters (which would require far more simulations). We encourage readers for which parameter uncertainty is relevant to implement system-specific simulations to investigate algorithm sensitivity.

| **Process** | **Component** | **Distribution** | **Parameter** | **Degree ($c$)** | **Value** |
|---|---|---|---|---|---|
| | | | | 1.0 | -0.010 |
| | | | | 1.5 | -0.015 |
| | | | | 2.0 | -0.020 |
| | | | $\gamma$ | 0.1 | 75 |
| | | | | 0.5 | 375 |
| | | | | 1.0 | 750 |
| | | | | 1.5 | 1125 |
| | | | | 2.0 | 1500 |
| **Archival** | Depth measurement $\left(y_t^{(D)}\right)$ | Uniform | $a_{y^{(D)}}$ | 1.0 | $b(\boldsymbol{s}_i) - 5$ |
| | | | $b_{y^{(D)}}$ | 1.0 | $b(\boldsymbol{s}_i) + 5$ |
